# Craters on the melanoma surface facilitate tumor-immune interactions and demonstrate pathologic response to checkpoint blockade in humans

**DOI:** 10.1101/2024.09.18.613595

**Authors:** Aya Ludin, Georgia L. Stirtz, Asaf Tal, Ajit J. Nirmal, Naomi Besson, Stephanie M. Jones, Kathleen L. Pfaff, Michael Manos, Sophia Liu, Irving Barrera, Qiyu Gong, Cecilia Pessoa Rodrigues, Aditi Sahu, Elizabeth Jerison, Joao V. Alessi, Biagio Ricciuti, Douglas S. Richardson, Jodi D. Weiss, Hadley M. Moreau, Meredith E. Stanhope, Alexander B. Afeyan, James Sefton, Wyatt D. McCall, Emily Formato, Song Yang, Yi Zhou, David P. Hoytema van Konijnenburg, Hannah L. Cole, Miguel Cordova, Liang Deng, Milind Rajadhyaksha, Stephen R. Quake, Mark M. Awad, Fei Chen, Peter K. Sorger, F. Stephen Hodi, Scott J. Rodig, George F. Murphy, Leonard I. Zon

**Author notes:** These authors contributed equally (Aya Ludin, Georgia L. Stirtz).

## Abstract

Immunotherapy leads to cancer eradication despite the tumor’s immunosuppressive environment. Here, we used extended long-term in-vivo imaging and high-resolution spatial transcriptomics of endogenous melanoma in zebrafish, and multiplex imaging of human melanoma, to identify domains that facilitate immune response during immunotherapy. We identified crater-shaped pockets at the margins of zebrafish and human melanoma, rich with beta-2 microglobulin (B2M) and antigen recognition molecules. The craters harbor the highest density of CD8^+^ T cells in the tumor. In zebrafish, CD8^+^ T cells formed prolonged interactions with melanoma cells within craters, characteristic of antigen recognition. Following immunostimulatory treatment, the craters enlarged and became the major site of activated CD8^+^ T cell accumulation and tumor killing that was B2M dependent. In humans, craters predicted immune response to ICB therapy, showing response better than high T cell infiltration. This marks craters as potential new diagnostic tool for immunotherapy success and targets to enhance ICB response.

## Introduction

Immune check-point blockade (ICB) cancer therapy, aimed at inducing CD8^+^ T cell responses against tumors, has significantly increased patients’ life expectancy and survival^1,2^. Mechanistic knowledge as to how CD8^+^ T cell responses are regulated and propagated within tumors is limited. However, recent reports of recurrent colocalization of T and myeloid cells^3–5^, suggest that the nature of immune response within tumors is regional rather than diffusive. How such regionality propagates anti-tumor immune responses, and how it is spatially assembled, organized, and regulated by the tumor is yet unknown. To date, a 3-dimensional, dynamic view of tumor-immune interactions *in vivo* has been challenging, limiting our understanding of the nature of anti-tumor immune response as it evolves within an intact tissue microenvironment. Such understanding is needed not only to improve treatment outcome, but also to accurately identify effective response to ICB treatment, as there is currently no pathological marker of ICB success other than tumor necrosis and fibrosis, and the presence of immune infiltration.

We have previously shown that human melanoma driver mutations, such as BRAF(V600E), can induce endogenous melanoma growth in zebrafish^6^. These tumors exhibit shared genomic and morphological characteristics of human melanoma^6–8^ and can be generated in immune-competent zebrafish with a highly conserved immune system^9^. The zebrafish model provides a unique opportunity to perform three-dimensional (3D) live imaging of immune cells as they interact in situ with endogenous tumors.

Here, we created a *cd8α*:EGFP transgenic fish and adapted a water flow system^10^ to allow live 3D time-lapse confocal imaging for over 15 hours. This enabled us to follow infiltrating CD8^+^ T cells as they move through the 3D architecture of endogenous tumors. Moreover, the extended length of imaging enabled us to identify very slow or rare processes undetected by previously reported live imaging studies. We found that rather than diffusely patrolling the tumor surface, CD8^+^ T cells aggregate in crater-shaped pockets on the melanoma perimeter, forming prolonged interactions with melanoma cells. We found the craters to be sites of CD8^+^ T cell antigen recognition where melanocytes lining the crater locally present elevated beta-2 microglobulin (B2M) levels and retain the CD8^+^ T cells in them for many hours. Craters exist in both zebrafish and human melanomas and become major sites of B2M-dependent tumor killing upon immune-stimulatory CpG ODN therapy in zebrafish. Highly multiplexed imaging of human melanoma shows similar MHC-high craters which preferentially harbor CD8^+^ T cells and PD-L1 high, CD163^+^ DCs. CD8^+^ T cell accumulation specifically within craters was found in patients responding favorably to ICB treatment, while patients with progressive disease post-treatment demonstrated disseminated CD8^+^ T cells infiltration throughout the tumor parenchyma. Furthermore, crater density in tumors was significantly higher in patients responding to treatment compared to non-responders, even when high levels of CD8^+^ T cells infiltration was detected in non-responders. Our study thus identifies physical regions, craters, on the tumor surfaces (internal and external tumor-stromal interfaces) in which the tumor is highly recognizable to CD8^+^ T cells. These regions promote and propagate an anti-tumor immune response upon treatment and represent a novel histopathological biomarker for the anti-tumor immune response elicited by ICB therapy.

## Results

### Generation of CD8^+^ T cell reporter zebrafish

To observe CD8^+^ cell interactions with melanoma in vivo, we created a transgenic zebrafish line that labels cells expressing *cd8a*. We cloned a region upstream of the *cd8a* zebrafish gene that contains two open chromatin areas identified by ATAC-seq analysis of sorted lck^+^ T cells (Figure S1A, top panel). The cloned fragment was used to drive cytoplasmic EGFP expression in *cd8a* expressing cells. EGFP expression was evident in the zebrafish thymus as early as 1-week post-fertilization and was prominent in juvenile fish thymus at 6 weeks post fertilization (Figure S1A, bottom panel). EGFP labeled a distinct population of lck^+^ cells in the fish major lymphoid organs (Figure S1B) that were round and approximately 10 µm in diameter (Figure S1B, top right image). RNA-seq of sorted EGFP^+^ cells from the thymus of a juvenile fish showed a gene expression profile characteristic of CD8^+^ T cells (Figure S1C), including expression of *cd8a*, *cd8b*, pan-T cell markers (*lck*, *cd3eap*), TCR signaling molecules (*zap70*), and transcription factors required for CD8^+^ T cell development (*gata3*, *runx3*). These cells did not express *zbtb7b*, which is required for CD4^+^ T cell development^11^. CD8^+^/CD4^+^ double-positive T cells exist in the thymi of juvenile fish and may contribute to the low levels of CD4 expression that was detected in this bulk population^9^. Taken together, the cd8a:EGFP transgenic line can be used to study CD8^+^ T cells and visualize their dynamics in vivo.

In addition to CD8^+^ T cells, the cd8α:EGFP fish labeled a population of myeloid cells (Figure S1D). These cells were 50-70 µm in diameter and had a distinct dendritic shape (Figure S1D,G). RNA-seq of sorted CD8^+^ myeloid cells from a melanoma revealed gene expression resembling murine CD8^+^ DCs. These CD8^+^ cells expressed the zebrafish myeloid markers *mpeg* and *mfap4*, but not the neutrophil marker *mpx* (Figure S1E). Transcription factors important for CD8^+^ DC development (*id2*, *batf3*, and *irf8*, but not *irf4*)^12^ or marking classical DCs (*zbtb46*)^13^ were highly expressed (Figure S1E,F). *cd8a*, but not *cd8b*, was expressed in this population (Figure S1F), characteristic of CD8^+^ DCs^14^. The pan-T cell marker *lck* was not expressed (Figure S1F), excluding T cell contamination. The CD8^+^ dendritic cells (DCs) internalized and presented AlexaFluor 594-conjugated ovalbumin in situ when injected intra-peritoneally (I.P), demonstrating an ability to uptake antigens (Figure S1G). This indicates that in addition to CD8^+^ T cells, the cd8α:EGFP transgenic fish marks a CD8^+^ DC population corresponding to murine CD8^+^ DCs^12,14^.

### T cells infiltrate melanoma tumors in zebrafish

We next visualized immune infiltration into zebrafish melanomas. We generated BRAF^V600E^/p53^null^ melanomas using two approaches. The first approach introduces an *mitfa* minigene, necessary for melanocyte formation, using the MiniCoopR vector injected into (BRAF^V600E^/p53^null^/nacre^null^) transgenic embryos that have been widely used in our lab^6,7^. The second system introduces de-novo the BRAF^V600E^ mutated protein expression, along with melanocyte specific CRISPR of p53 and tyrosinase to create non-pigmented tumors^15^. This approach produces tumors in lower frequency but enables us to generate tumors in any zebrafish line. We found by single-cell RNA-seq (scRNA-seq) that lck^+^ T cells infiltrating zebrafish tumors (Figure S2A,B) contain CD8^+^ T cells and CD4^+^ T cells, including Foxp3^+^ regulatory T cells (Figure S2C,D). To assess T cell dynamics in vivo, we first performed longitudinal still imaging of an individual zebrafish and visualized T cell infiltration as the tumor developed (Figure 1A). Early melanocytic (pre-tumorous) lesions grow radially beneath the fish scale. At this stage, lck^+^ T cells, with very few CD8^+^ cells, retained their normal distribution along the scale edges, creating tessellated structures^16^ (Figure 1A, left). These tessellated structures along the fish scale were reported to serve as peripheral immune surveillance sites that functionally resemble lymph nodes^16^. As the tumor protrudes from the scales to form a mass, the tessellated pattern is disrupted (Figure 1A, middle) and ultimately disappears, while multiple CD8^+^ T cells infiltrate the tumor (Figure 1A, right). At this stage, we noted that CD8^+^ T cells infiltrating the tumor are enriched in areas devoid of mitfa^+^ melanocytes (Figure 1A, right enlarged), suggesting that CD8^+^ T cells infiltrating tumors may aggregate at specific domains on the tumor margins.

**Figure 1.**
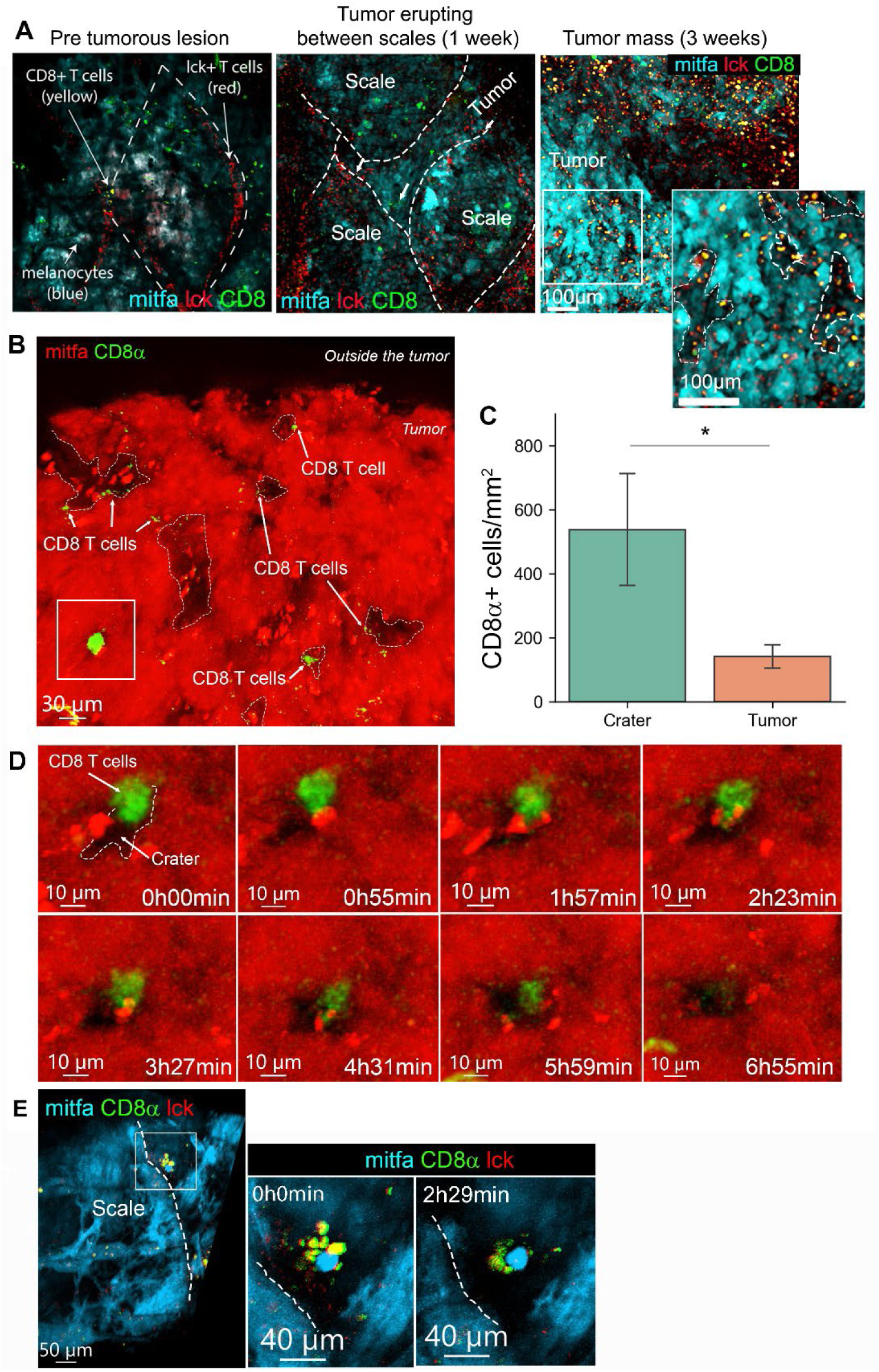
CD8^+^ T cells are preferentially retained in craters, interacting with tumor cells. (**A**) Longitudinal imaging of a mitfa:BFP tumor across stages of development in a (cd8a:EGFP;lck:mCherry) transgenic fish. Left: Radial growth of melanoma below the scale. T cell skin distribution remains normal. Middle: Same tumor after one week. Slightly protruding melanoma and the tessellated T cell structure is disrupted. Right: Same tumor after three weeks from initial imaging. A protruding tumor, where CD8^+^ T cells infiltration is evident. Enlarged-CD8^+^ T cells are found in areas of low mitfa expression (marked by dashed white lines). (**B**) 3D projection of a melanoma tumor surface in a cd8α:EGFP transgenic zebrafish (representative of over 200 tumors imaged in this study). Arrows mark CD8^+^ T cells, which are found in mitfa:mCherry devoid pockets, i.e. craters. (**C**) CD8^+^ cell density (cells per mm^2^) in crater vs. tumor area (n=5 fish. Data is mean±SE, Mann Witney U test, *p value=0.015). (**D**) Snapshots from time lapse imaging of the area marked by the rectangle in (B). Time is hours:minutes. A CD8^+^ cell interacting with a melanoma cell over a crater, then enters the crater after 6 hours. (**E**) Snapshots from time lapse imaging of a scale edge crater, show multiple CD8^+^/lck^+^ T cells interacting with a melanoma cell in a crater. Dashed line marks the edge of the scale. Time is hours:minutes.

### CD8^+^ T cells localize to craters and engage with tumor cells at the melanoma surface

To study this phenomenon, we quantitatively analyzed 3D confocal images of endogenous zebrafish melanoma. We found that CD8α^+^ cells preferentially localized to areas devoid of melanocytes that form crater-shaped pockets lined by melanoma cells (Figure 1B), here termed “craters”. This was evident in all melanoma tumors imaged during this study (>100 fish). We developed an algorithm to segment the craters, allowing determination of crater size (materials and methods, Figure S3A) and its interaction with other objects, including CD8^+^ T cells (Figure S3B). We found that the craters had an average diameter of 50 µm (ranging between 20 µm to 150 µm) (Figure S3C) and constitute about 12% of the surface of an untreated tumor (Figure S3D). CD8^+^ T cell density was higher in craters compared to the rest of the tumor in a statistically significant manner (Figure 1C). Moreover, we found that the majority of CD8^+^ T cells were within 5 µm of a crater and their numbers decreased as the distance to the nearest crater grew (Figure S3E), suggesting that CD8^+^ T cells preferentially localize to craters. To visualize CD8^+^ cell activity within the craters, we adapted a water flow system to keep zebrafish alive and anesthetized under a confocal microscope^10^ (see materials and methods). We used this system to track 3D cellular dynamics for 15-20 hours via time-lapse imaging. Within the tumor mass, CD8^+^ T cells lingered in craters and engaged with melanoma cells for hours, then observed entering the crater (Figure 1D, video S1), or moving out of the crater (Figure S4A, video S2). At the scale edge of the tumor, which are sites of immune surveillance^16^, clusters of CD8^+^ T cells were found in craters (Figure 1E, S4B) directly engaging a tumor cell for many hours (Figure 1E, video S3) in a manner resembling tumor recognition and killing, as seen in vitro^17^.

To evaluate the craters proximity to tumor vasculature, we imaged tumors grown in *flk1*:EGFP and *fli1a*:dsRed transgenic zebrafish, marking blood vessels^18^. We found that craters emerge at the perivascular area, adjacent to blood vessels, closely associated with fli1a^+^/flk-vessels (Figure S4C). Using scRNA-seq of sorted fli1a^+^/flk^+^ and fli1a^+/^flk^-^ cells, we found that the fli1a+/flk-endothelial cells correspond to venous and lymphatic cells (Figure S5A-D), suggesting that venous and lymphatic vessels are associated with craters. CD8^+^ T cells in the craters were found contacting either fli1^+^ vessel (Figure S4D, area 1) or melanoma cells (Figure S4D, area 2). In addition to CD8^+^ T cells, craters harbor mpeg^+^ monocytes (Figure S4E, area 1). CD8α^+^/mpeg^+^ DCs, that are similar to CD8^+^ DCs (Figure S1D-G), that were shown in mammals to cross-present antigens with high efficiency to CD8^+^ T cells^14^, could be found over craters in untreated tumors (Figure S4E, area 2). Regarding extra cellular matrix components, using second harmonic generation^19^, we found fine collagen fibers extending into the craters (Figure S4F) from the capsule surrounding the zebrafish tumor (Figure S4F, S7G).

### The craters are sites of elevated B2M levels and antigen recognition for CD8^+^ T cells

CD8^+^ T cell behavior within craters strongly resembled T cell behavior reported to occur during antigen recognition, when T cells form sustained interactions with their recognized target cells^20^. To further characterize the crater areas on the melanoma surface, we performed high resolution spatial transcriptomics (Slide-seqV2)^21^ on zebrafish tumors. We identified pocket-shaped areas akin to craters, lined by cells expressing a melanoma gene signature, occasionally containing cells expressing a gene signature characteristic of T cells (Figure 2A). We then performed differential gene expression analysis (see materials and methods) between melanoma cells at the craters’ border and melanoma cells not associated with craters, at the tumor surface (as shown in Figure 2A). Melanoma cells in crater areas presented elevated expression of *b2m* (Figure 2B), encoding B2M, a core molecule of the MHC class I complex that enables tumor recognition by CD8^+^ T cells and its loss was shown to lead to immune evasion^22^. To examine whether B2M has a role in CD8^+^ T cell retention to craters, we designed a gRNA to perform CRISPR knockdown of B2M in zebrafish. The gRNA was inserted into a melanoma-specific CRISPR vector^23^ and injected into cd8α:EGFP transgenic embryos along with melanoma-generating vectors (described above) to induce tumors with melanoma-specific B2M depletion. Imaging and quantification of these tumors revealed that following B2M depletion in melanoma cells, a higher proportion of CD8^+^ T cells were found embedded in the tumor mass rather than aggregated within craters when compared to B2M intact, wild type (WT) tumors (Figure 2C), manifested in a significant reduction in the ratio between CD8^+^ T cells in craters vs. embedded in the tumor mass in B2M depleted tumors compared to WT control (Figure 2D). This indicates that impaired melanoma recognition leads to either reduced migration of CD8^+^ T cells to craters or decreased duration of CD8^+^ T cell dwelling within craters. The overall CD8^+^ T cell number was not significantly changed compared to B2M intact tumors (Figure S6A), nor was crater coverage (Figure S6B). Taken together, loss of B2M, which hampers MHC class I assembly in melanoma cells^22^, reduced CD8^+^ T cell affinity to the craters. This, together with the characteristic long-term interactions of CD8^+^ T cells with melanoma in craters, indicates that craters are sites where CD8^+^ T cells recognize tumor cells and engage them.

**Figure 2.**
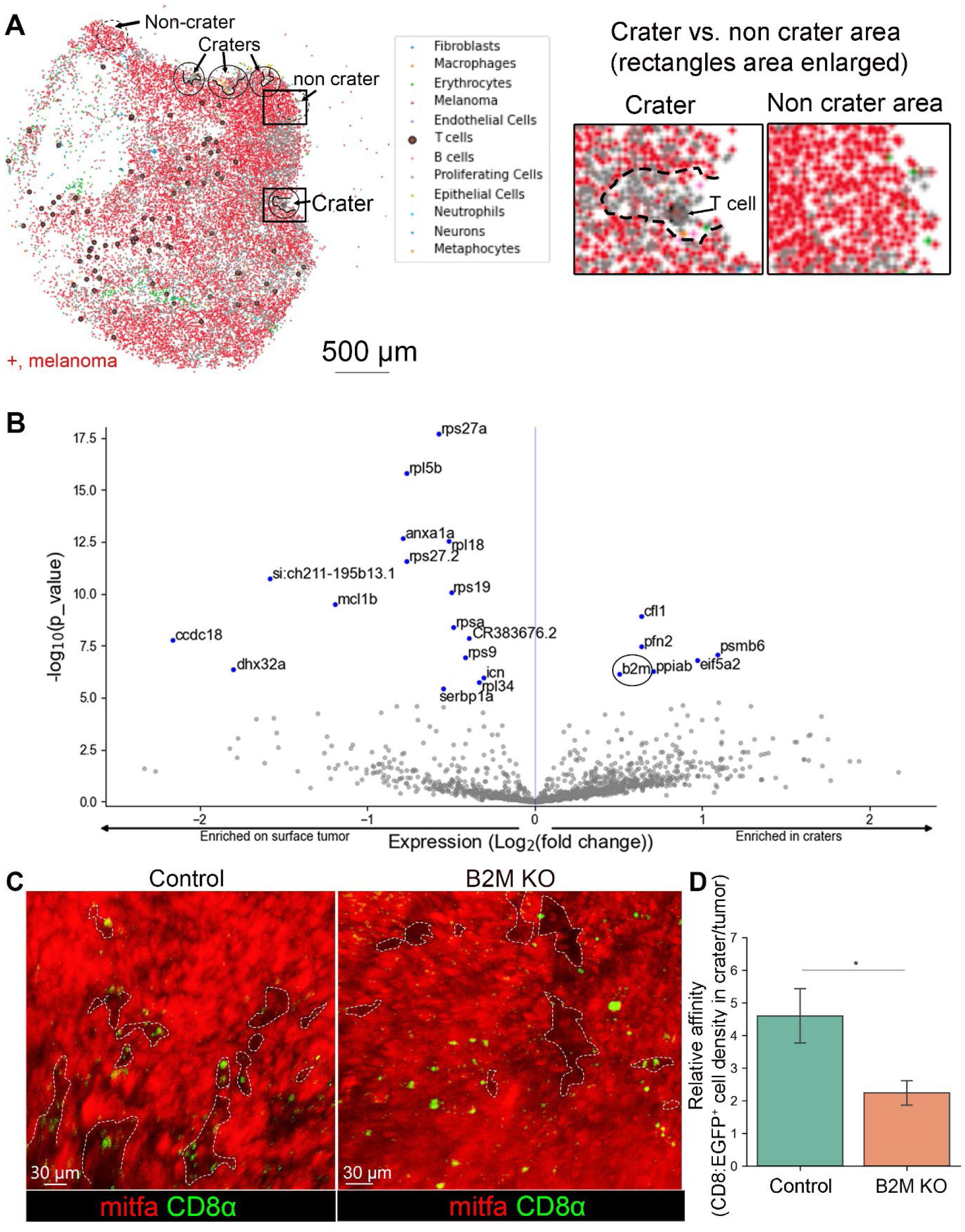
Craters are sites of elevated B2M expression which retains CD8^+^ T cells in them. (**A**) Left: Slide-seq gene expression map of zebrafish melanoma tumor depicting an array of cells including CD8^+^ T cells (large purple circle) and mitfa^+^ melanoma (red). Crater areas are circled in black, non-crater areas are circled in dashed black line. Right: enlarged view of the crater and non-crater areas marked by rectangles in the image on the left. (**B**) Volcano plot identifying differential gene expression in crater vs. non-crater associated tumor cell. B2M is circled. (**C**) 3D live imaging of control(intact) vs. B2M depleted mitfa:BFP tumors grown in (cd8α:EGFP/lck:mCherry) zebrafish (Mitfa color switched by Imaris to red for better visualization). Craters are marked with dashed lines. (**D**) The relative affinity of CD8^+^ T cells to craters, i.e CD8^+^ cell density in craters/non crater tumor surface, in control, intact B2M-and melanoma specific B2M-depleted tumors. (n=4 control, n=6 B2M CRISPR ablated. All CRISPR sequenced to verify mutation. Data is mean±SE, T test. *=p-value 0.02).

### Activated CD8^+^ cells accumulate in enlarged craters following immune stimulation

Antigen recognition is an essential step in CD8^+^ T cell activation^24^ required for anti-tumor cytotoxicity, suggesting that craters can facilitate tumor killing during treatment. To test whether craters can become sites of effective T cell-mediated tumor killing, we treated tumor bearing zebrafish with immunotherapy. The identification of the homologous PD-1, PD-L1 and CTLA-4 genes in zebrafish are still ongoing^25,26^, making ICB treatment in zebrafish challenging. However, CpG ODN, a TLR agonist in clinical trials for melanoma as an immunotherapy^27^, has been shown to induce Interferon-γ (IFN-γ) expression in zebrafish^28^. We therefore treated tumor-bearing zebrafish with daily intra-tumoral injections of CpG ODN (Figure S7A) and monitored the treated fish 24 hours after each injection. We found that seven daily injections of CpG ODN, but not of PBS vehicle, caused visible shrinkage of tumors (Figure S7B, upper and middle panels). Following four daily injections of CpG ODN but not vehicle, tumors presented large craters containing many CD8^+^ T cells (Figure 3A). We also visualized disruption of the collagen layer surrounding the tumor mass (Figure S7G), consistent with active T cell trafficking and infiltration.

**Figure 3.**
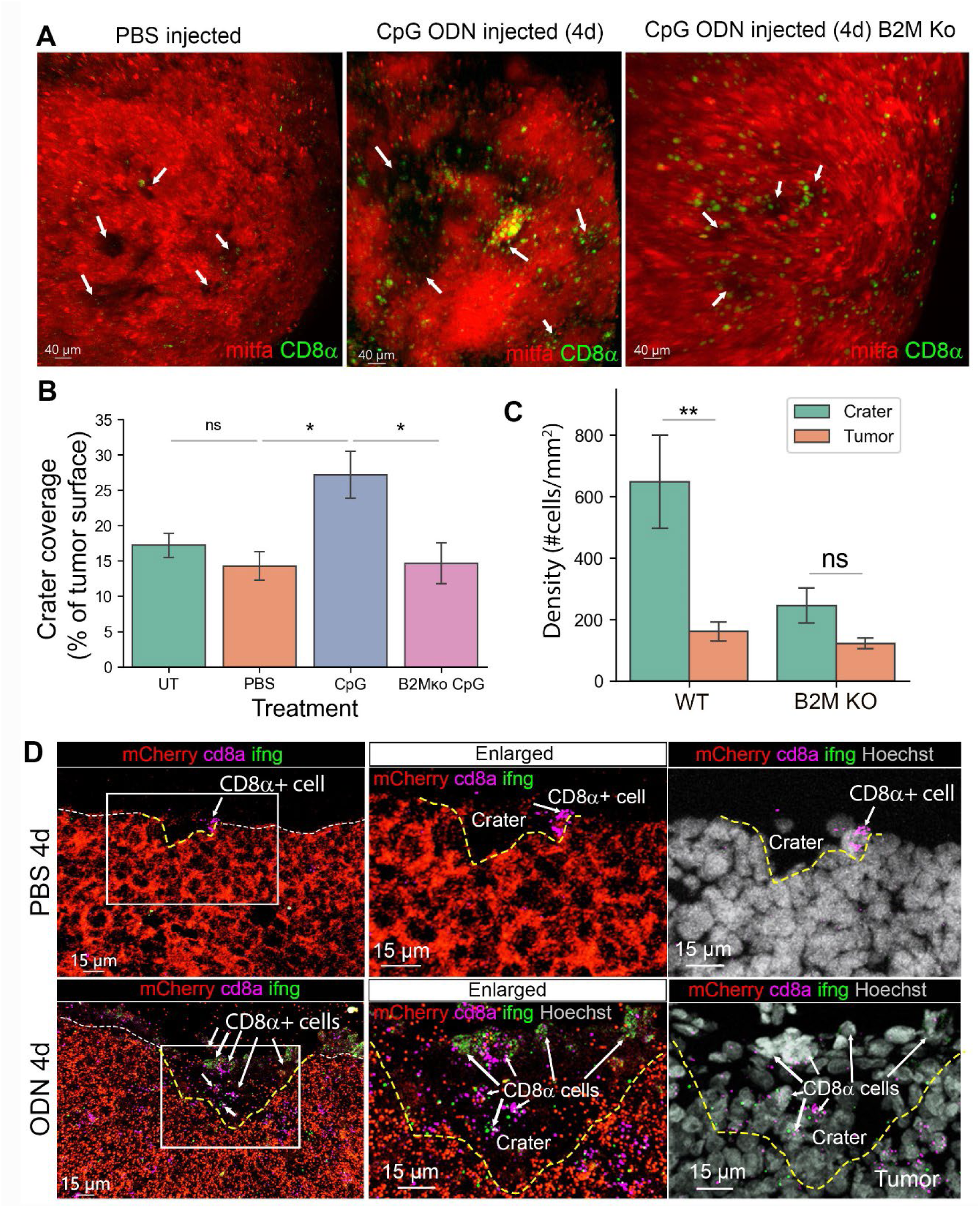
Activated CD8^+^ T cells aggregate in craters following immune stimulation by CpG ODN. **(A)** 3D live imaging of mitfa:mCherry melanoma tumors in cd8α:EGFP transgenic fish after 4 daily intra-tumoral injections of vehicle (PBS and water, referred here as PBS) or CpG ODN, 24h post last injection, injected into either B2M WT or KO tumors. Arrows indicate craters. **(B)** Crater coverage in untreated-, PBS-CpG ODN- and tumor specific B2M ko, CpG ODN injected tumors after 4 daily injections (n=UT-5, PBS-5, CpG ODN-5, B2M KO CpG ODN-4 fish. Data are mean±SE, Mann Whitney U test, *p value=0.031, ns=0.42). **(C)** CD8^+^ T cell density in craters vs non crater tumor surface in B2M intact (WT) and B2M tumor specific KO tumors, treated with 4 daily injections of CpG ODN (n=5 WT and n=4 B2M KO CpG ODN treated fish, Data is mean±SE, Mann Whitney U test, **p value =0.007). **(D)** RNA Scope for mCherry, cd8a and ifng RNA transcripts in sections from tumors treated with 4 daily injections of PBS (vehicle) or CpG ODN. Dashed white line marks the tumor surface. Yellow dashed line marks craters that are depressions in the tumor surface. Arrows indicate CD8^+^ cells (representatives of 3 PBS and 3 CpG ODN treated fish undergone RNA Scope analysis).

Quantification of the images showed a statistically significant increase in crater coverage (i.e. percent of tumor surface accounted for by craters) following CpG ODN treatment (Figure 3B). CD8^+^ T cell density remained higher within craters compared to the tumor mass (Figure 3C) and their relative affinity to the craters, i.e. the ratio of CD8^+^ T cells within craters/embedded in tumor remained high as in untreated tumors (Figure S7C), consistent with persistent spatial localization of CD8^+^ T cells to craters. We found these effects to be B2M-mediated. We injected CpG ODN daily into B2M KO tumors generated in the same fish strain as the above described B2M wild type. Unlike B2M WT tumors, CpG ODN injections for seven days did not induce shrinkage of the B2M KO tumors (Figure S7B, lower panel). Four daily injections of CpG ODN into B2M-depleted fish resulted in CD8^+^ T cells recruitment to the tumors despite the loss of B2M expression by the tumor (Figure S7D). However, unlike in B2M WT tumors, many CD8^+^ T cells did not come in contact with the tumor and remained spatially localized >5µm away from the tumor (Figure S7E,F). The CD8^+^ T cells that were in contact with the tumor were abundant in the tumor parenchyma, resulting in reduced CD8^+^ T cells density (Figure 3C) and thereby, reduced affinity (Figure S7C) of CD8^+^ T cells to craters in CpG ODN treated B2M KO tumors. There was no evident increase in crater volume following treatment when B2M was ablated from the tumor cells (Figure 3A, right). Indeed, crater coverage in B2M-depleted tumors treated with CpG ODN for four days remained unchanged and was similar to PBS or untreated wild type tumors (Figure 3B, S6C). The above indicates that B2M mediated the attachment of CD8^+^ T cells to the tumor and localization of infiltrating CD8^+^ T cells to craters.

We next examined the behavior and phenotype of the CD8^+^ T cells within craters following CpG ODN treatment. Live imaging showed multiple CD8^+^ T cells within the craters, interacting with multiple melanoma cells (video S4). Using RNA Scope to detect *ifng1* (encoding IFN-γ which is expressed by activated CD8^+^ T cell^29^), *cd8a,* and *mCherry* gene expression in CpG ODN- and vehicle-treated tumors, we found that CD8^+^ T cells associated with craters in vehicle-treated tumors were mostly *ifng1* negative (Figure 3D, upper panel). In CpG ODN-treated tumors, multiple CD8^+^ T cells expressing *ifng1* were visible in enlarged craters, suggesting that CD8^+^ T cells in the craters were activated.

As craters are sites of tumor recognition and engagement by CD8^+^ T cells, aggregation of activated CD8^+^ T cells within craters indicates that the craters may serve as primary sites of the anti-tumor immune response.

### Craters are sites of tumor killing following immune stimulation

The observation of increased crater coverage together with the identification of activated CD8^+^ T cells within craters of immunotherapy-treated tumors when B2M is intact suggests that CD8^+^ T cells kill tumor cells within the craters in a B2M dependent manner. To assess this, we performed whole-mount TUNEL assay, enabling a 3D view of the tumor surface, on tumors following four daily injections of CpG ODN or vehicle into B2M intact or depleted tumors. We found that in vehicle-treated tumors and CpG ODN treated B2M KO tumor, very few apoptotic TUNEL^+^ cells were found on the tumor surface (Figures 4A left and right panels, S7H. There were 176 TUNEL^+^ cells in PBS vs, 1738 cells following CpG ODN treatment in a B2M wild type (WT) tumor and 460 TUNEL^+^ cells in CpG ODN treated B2M KO tumor). Following four daily injections of CpG ODN, TUNEL^+^ cells were detected preferentially within and around the rim of craters in CpG ODN treated B2M WT tumors but not in PBS treated WT tumors or CpG ODN treated B2M KO tumors (Figures 4A,B, S7H). CD8^+^ T cells could be found in direct contact with TUNEL^+^ melanoma cells (Figure S7H, left, enlarged), an association consistent with T cell-mediated cytotoxicity. Taken together, these results indicate that craters are a major site of tumor killing upon treatment, mediated by B2M.

**Figure 4.**
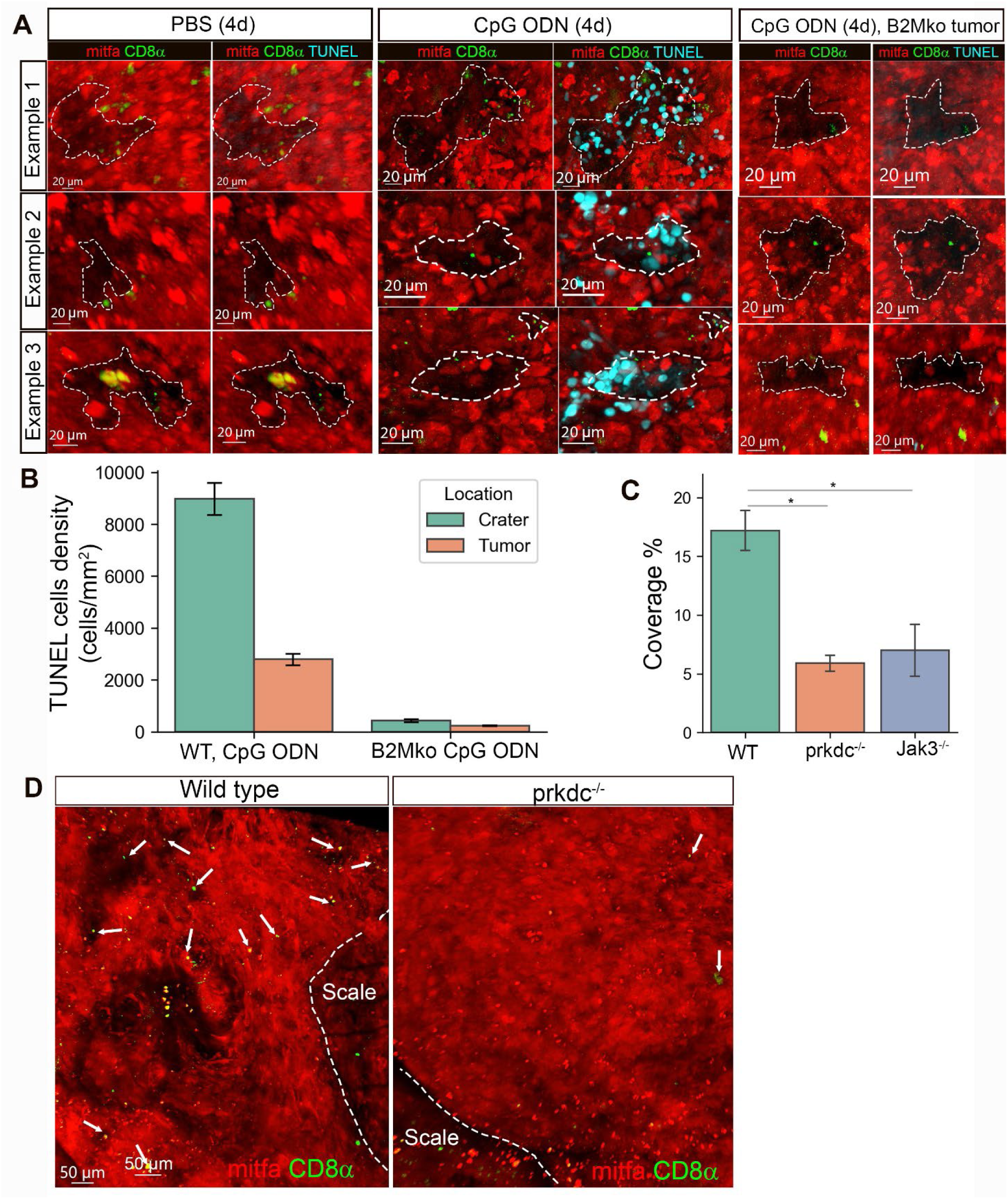
Craters are sites of tumor killing and are remodeled by T cells. **(A)** 3D confocal images of whole mount TUNEL assay in tumors undergone 4 daily intratumorally injections with PBS or CpG ODN, showing examples of 3 craters in either PBS- or CpG ODN- or B2M KO tumor, CpG ODN treated tumors, each representative of 2 PBS and 2 CpG ODN treated fish. (**B**) TUNEL^+^ cells density in craters and non-crater tumor area (calculated using bootstrap for TUNEL^+^ cells. CpG ODN treated sample contained 1738 TUNEL^+^ cells. PBS-176 TUNEL^+^cells, CpG ODN treated B2M KO tumor-460 TUNEL^+^ cells) (**C**) Crater coverage in mitfa:mCherry tumors of cd8α:EGFP (immunocompetent wild type), cd8α:EGFP;prkdc^-/-^ or jak3^-/-^ transgenic fish (n= 5 wt, 2 prkdc^-/-^, 3 jak3^-/-^ fish. Mean± SD, T-test,*p-value (cd8a:EGFP;prkdc^-/-^)=0.011, *p value jak3^-/-^=0.01) (**D**) Representative 3D confocal images of mitfa:mCherry tumors in wild type cd8a:EGFP and (cd8a:EGFP;prkdc^-/-^) transgenic fish. Arrows indicate CD8^+^ T cells.

These results further indicated that CD8^+^ T cells contribute to sculpting the crater architecture, by tumor killing. To test whether T cells are involved with crater formation, we compared wild type (immunocompetent) tumors to tumors from two distinct T cell-depleted zebrafish lines: cd8α:EGFP;prkdc^(-/-)^ transgenic zebrafish lacking T cells and B cells^30^ and jak3^(-/-)^ transgenic zebrafish lacking T cells and putative NK cells^30^. We found that crater coverage in tumors of both T cell-deficient strains was significantly lower, but not absent, compared to tumors of T cell-competent fish (Figure 4C,D), indicating that T cells determine crater coverage in tumors and suggests that the craters may originate from topographical changes in the tumor that are sculptured and maintained by T cells. Taken together, we found that during immunostimulatory treatment, CD8^+^ T cells localize to craters and exert their anti-tumor immune response within the crater. These effects are mediated by B2M expression.

Last, we found evidence that CD8^+^ myeloid corresponding to CD8^+^ DCs (Figure S1E-G), that may be found in craters in untreated tumors (Figure S4E, area 2), can increase in number following immune stimulatory treatment. TGF-β is a broad immunosuppressive molecule capable of inhibiting antigen presentation by DCs^31^. Drugs aimed to inhibit TGF-β are currently in clinical trials^31^. We inhibited TGF-β for 24 hours using the TGF-β inhibitor (TGF-βi) SB431542 and found increased accumulation of CD8^+^ cells (Figure S8A,B,C), including CD8α^+^/mpeg^+^ cells (Figure S8A, example 2) in craters, suggesting that antigen presenting cells may accumulate in craters as well upon treatment. Using long-term time-lapse imaging of TGF-βi treated tumor, we saw a lengthy interaction of CD8^+^ T cells with a melanoma cell at the scale edge, followed by its delivery to a CD8^+^/mpeg^+^ dendritic shaped cell and uptake of an mCherry fragment by the CD8^+^ DC (Figure S8D,E, video S5), suggestive of the initial steps of antigen uptake, captured live following TGF-β inhibition. Taken together, results suggests that immune stimulation can induce both accumulation of activated CD8^+^ T cells and antigen presenting cells in craters, making them sites of active tumor killing.

### Human melanoma tumors exhibit craters harboring densely packed CD8^+^ T cells

Using the zebrafish model, we learned that CD8^+^ T cells localize to crater-shaped immune regulatory sites in the tumor, creating spatially defined niches for tumor antigen recognition and killing by CD8^+^ T cells. To determine if such craters exist in human melanoma, we studied a total of 20 primary human melanoma tumors and 24 melanoma metastases from untreated patients, using Cyclic Immunofluorescence (CycIF) and Akoya Opal multiplexed immunofluorescence (mIF) (Figure 5B, Table 1). We found that all examined tumors harboring CD8^+^ T cells, both primary melanoma and melanoma metastases, contained craters. In human melanoma, the craters are very small pockets lined up by melanoma cells breaching into the tumor mass from stromal layers, at either the tumor border with the adjacent tissue (Figure S9A), or at the perivascular area, which is the stromal layer that encircles blood vessels (Figure 5A, illustrated in 5C). These regions have been shown to contain DCs and T cells in mice^32^ and human^33^. At the perivascular area, craters breach out as small (∼50 µm), pockets from the margins of the αSMA and collagen rich layer surrounding the blood vessels into the tumor mass (Figure 5A, S9B, illustrated in 5C). Similar craters extended from the margins of the dense fibrous, αSMA^+^ layer that separates the tumor from adjacent tissue (Figure S9A). As in zebrafish, fine collagen fibers extended into the craters from the collagen layer of the perivascular area (Figure S9B). This feature helps distinguish craters from the perivascular area, in a pathological examination of human samples. The craters characteristically contained CD8^+^ T cells and CD163^+^ dendritic cells in close proximity and were lined up by melanoma cells throughout-sides and bottom of the crater (Figure 5A, S9A). To calculate cell density and define the crater cellular composition in human tumors, we used cell segmentation followed by cell clustering of CyCIF data (see Materials and methods, Figure S10A,B) that enabled us to characterize the cells within the craters (Figure S10C). We found that as in zebrafish, CD8^+^ T cell density was the highest within craters (Figure 5D), compared to either the rest of the tumor margins or embedded within the tumor mass (areas defined as demonstrated in Figure S10A), indicating that CD8^+^ T cells cluster within craters. When examined the crater contents, we found that the craters mostly harbor CD8^+^ T cells and CD163^+^/CD11c^+^ DCs, (Figure 5E,F). Cytotoxic CD8^+^ T cells (CTL) expressing Granzyme B (GzmB) could be found in craters, in lower numbers than GzmB^-^/CD8^+^ T cells (Figure 5E,F). CD4^+^ T cells were mostly found outside the craters (Figure 5E,F). CD4^+^/ Foxp3^+^ T regulatory cells were found embedded in the tumor mass or in the perivascular area, occasionally close to the crater edges but rarely inside (Figure 5E,F). GzmB^+^ cells that were not T cells were also observed in human melanoma tumors but were rarely found within craters (Figure 5E,F).

**Figure 5.**
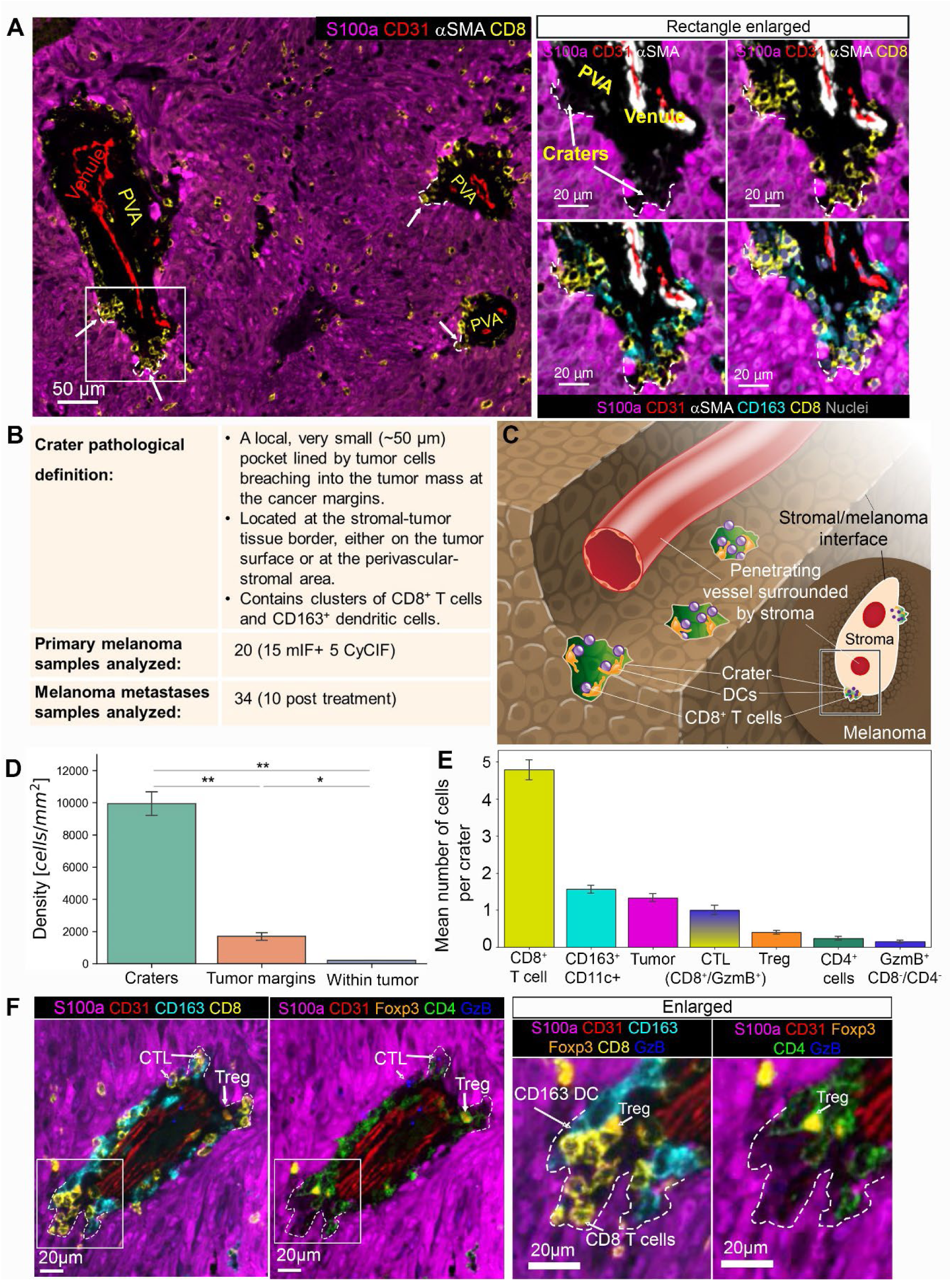
Craters are found in human melanoma, breaching from the tumor margins. **(A)** Left: low magnification of a representative human primary nodular melanoma (CyCIF). Craters are marked by arrows and white dashed lines. Right: the area marked by the rectangle in the left image is enlarged, showing CD8^+^ T cells and CD163^+^ DCs packed together within craters (marked by arrows and white dashed lines). **(B)** A table summarizing the craters features and samples used in this study. **(C)** a graphical illustration of craters (colored green) breaching from a perivascular space into the melanoma tumor. The content of the perivascular area was omitted for clear view of the craters. **(D)** CD8^+^ T cell density in craters, tumor margins and embedded in the S100a^+^ tumor mass (within tumor). (n=2 tumors. Data is mean± SE, T-Test, ** p value= crater/border=0.008, crater/tumor=0.005, *p value=0.023). **(E)** Cellular composition of craters showing mean number of cells per crater for each cell type (n=714 craters, Mean± SE). **(F)** A representative crater image, showing the location of CD8^+^ T cells, CD163^+^ DCs, CD4^+^ cells, CD4^+^/FOXP3^+^ T regulatory cells, and CD8^+^/GzmB^+^ cytotoxic T cells at the crater area. Abbreviations: CTL=cytotoxic T cells, Treg=T regulatory cells, GzmB= granzyme B. DCs= CD11c+ dendritic cells.

When examined crater formation in different stages of tumor development, we found that craters were only documented in the vertical (tumorigenic) phase of invasive growth (Figure S11C,D) and not in the earlier, more superficial radial phase of primary melanoma growth (Figure S11A,B). This is consistent with the known prognostic significance of the presence and extent of T cells that infiltrate the tumor-stromal interface of melanomas in the vertical/nodular stage of their growth^33,34^. Thus, craters are associated with the vertical growth phase of melanoma which represents a transition from non-metastasizing radial growth to invasive vertical growth, and in which the presence of tumor infiltrating lymphocytes (TILs) is a recognized prognostic attribute.

### Craters in human melanoma are sites of antigen presentation, rich with PD-L1

In zebrafish, we identified high expression of B2M in craters, which is a key component of MHC class I. In human melanoma, we found increased expression of HLA-A, corresponding to MHC class I, in the S100a^+^ tumor cells lining the craters (Figure 6A,B). We also noted that high levels of HLA-A are seen in cells inside the craters. We found that CD163^+^/CD11c^+^ DCs within craters present a statistically significant increase in the levels of the major histocompatibility complexes HLA-A and HLADPB1, corresponding to MHC class I and II respectively, compared to elsewhere in the tumor (Figure 6C,D). We further identified significantly elevated levels of MART1 melanoma antigen staining in CD163^+^/CD11c^+^ DCs within the craters (Figure 6C,D), suggesting that dendritic cells within craters take up and present melanoma antigens. In addition, PD-L1 expression was significantly higher in CD163^+^/CD11c^+^ DCs within the craters compared to the rest of the tumor (Figure 6E, F, Figure S12A), marking the craters as pockets rich with PD-L1 (Figure 6F, upper panel). This is consistent with previous reports of PD-L1 being expressed primarily by myeloid cells, DCs, and macrophages at the melanoma margins^33^. Of note, PD-L1 staining was significantly higher in CD163^+^ DCs contacting the CD8^+^ T cells in the craters compared to elsewhere in the tumor, even when comparing CD163^+^ cells at the same perivascular area (Figure 6F, lower panel, orange vs. blue arrows), suggesting that CD8^+^ T cells are exposed to not only antigens but also to PD-L1 within the crater. CD8^+^ T cells within and outside of the craters expressed markers of T cell dysfunction (Figure S12B), including PD-1. This suggests that CD8^+^ T cell dysfunction is not determined in the crater. However, it indicated that CD8^+^ T cells can respond to PD-L1, as they express its receptor PD-1 (Figure S12B). As in zebrafish, we found that crater formation correlates with the presence of CD8^+^ T cells in untreated tumors. When reviewing untreated, lowly infiltrated, cold tumors (eight primary melanoma and nine metastases) and untreated highly infiltrated, hot tumors (seven primary melanoma and nine metastases), we found that the perivascular areas in cold tumors exhibited few to no craters, while multiple craters were found in highly infiltrated, hot, tumors (Figure 6G).

**Figure 6.**
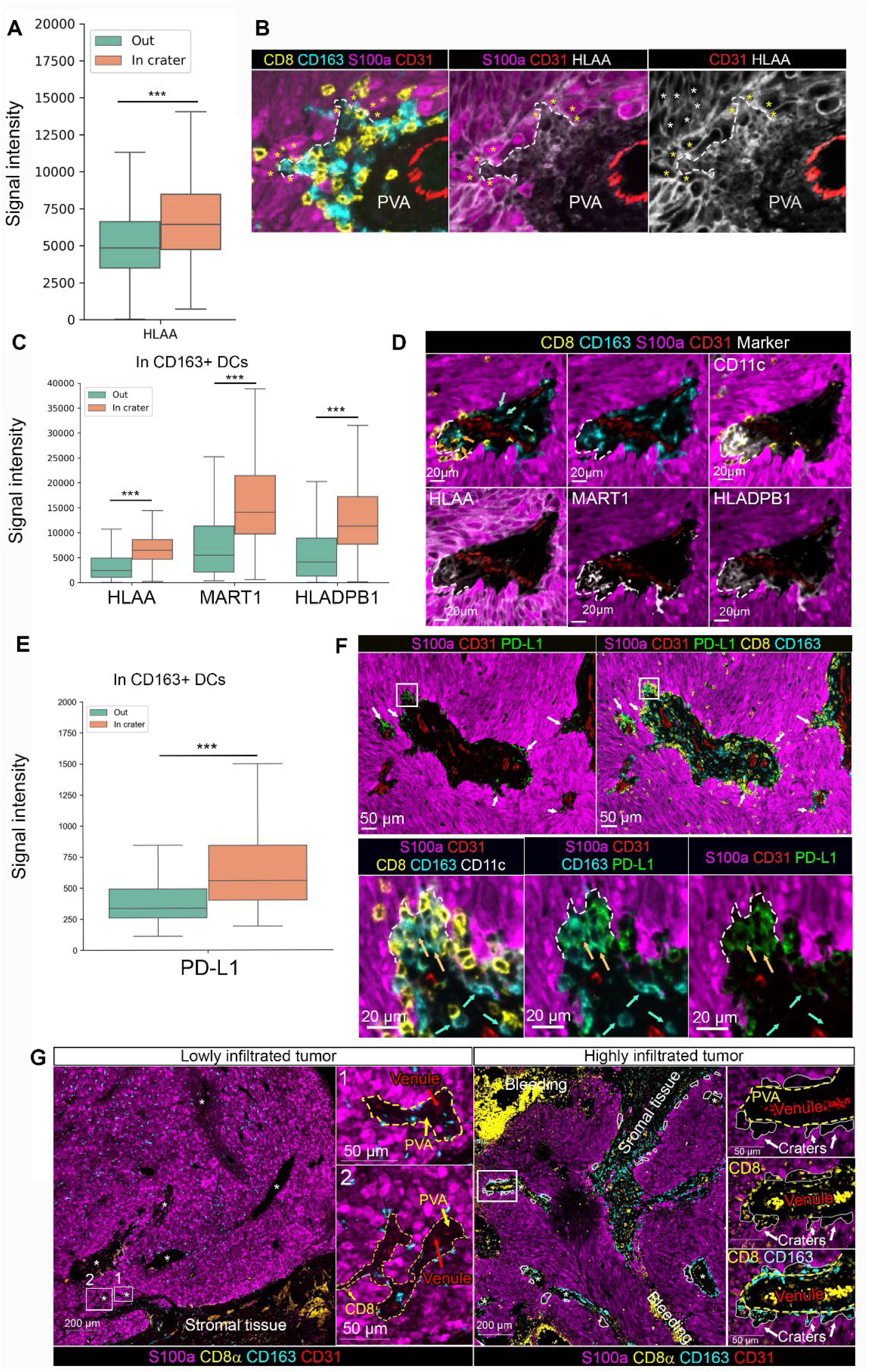
Craters in human melanoma are sites of high antigen presentation molecules and PD-L1 expression. (**A**) Boxplots plotting signal intensity distribution of HLA-A staining in S100a melanoma cells in craters compared to elsewhere in the tumor (termed, “Out”) (n=2 tumors, 714 craters, ***p value<0.001) **(B)** Representative image of a crater (marked by dashed white line), shown for S100a, CD8 and CD163 staining (right), HLA-A and S100a staining and HLA-A staining only to view the differentially high signal intensity in the melanoma cells lining the crater (marked by yellow asterisks) compared to the adjacent tumor cells outside the crater (white asterisks, marking a few examples). **(C)** Boxplots plotting signal intensity distribution of HLA-A, MART1 and HLADPB1 staining in CD163^+^/CD11c^+^ DCs in craters compared to elsewhere in the tumor (termed, “Out”) (n=2 tumors, 714 craters, ***p value<0.001) (**D**) Representative image of a large crater breaching out of the margins of a perivascular area, containing CD163^+^ DCs stained for HLA-A, MART1 and HLADPB1. (**E**) Boxplots plotting signal intensity distribution of PD-L1 staining in CD163^+^/CD11c^+^ DCs in craters compared to elsewhere in the tumor (“Out”) (n=2 tumors, 714 craters., ***p value<0.001). (**F**) Upper panel: low magnification of a tumor area showing craters as pockets with high levels of PD-L1 within them (white arrows), breaching into the tumor from the perivascular area (PVA). Lower panel: area marked by the rectangle is enlarged to show a crater (marked by dashed line) containing PD-L1 expressing CD163^+^/CD11c^+^ DCs. Blue arrows indicate CD163^+^/CD11c^+^ DCs (with low PD-L1 staining) outside the crater. Orange arrows indicate CD163^+^CD11c^+^ (PD-L1^+^) DCs inside the crater. **(G)** multiplex imaging of lowly and highly infiltrated tumors, showing lack of craters (marked by white lines) stemming from the peri-vascular areas (PVA, marked with dashed yellow line) in lowly infiltrated tumors compared to highly infiltrated tumors. Areas at the rectangles in the low magnification images to the left are enlarged at the right of each image.

### Craters demonstrate an effective anti-tumor immune response during ICB treatment regardless of CD8^+^ T cells infiltration levels

The current standard for pathologists to evaluate success with ICB is to examine the diffuse infiltration of tumor parenchyma with CD8^+^ T cells and assessment of fibrosis and necrosis. We next explored if craters could provide a more accurate assessment of ICB therapy success in patients. In zebrafish, craters harbor an effective anti-tumor immune response following T cell activation that leads to tumor killing. We examined residual metastases left following ICB therapy. Unlike samples from patients who did not respond to treatment, samples from patients who were declared clinically responsive to treatment are uncommon and difficult to obtain, as partial and complete responders may not require additional surgery. Nevertheless, we were able to study seven cases clinically presenting with progressive disease (PD), termed here “non-responders”, and three cases of durably responding patients, i.e. two cases presenting a stable disease (SD) and one complete response (CR), termed here “responders”. All patients were treated with anti-PD-1 and anti-CTLA4 antibodies. We found that craters were significantly more frequent along the stromal-tumor interface, termed here “tumor margin” (Figure 7B), in samples of responders compared to non-responders (Figure 7A). Thus, in these specimens, craters correlated with ICB success.

**Figure 7.**
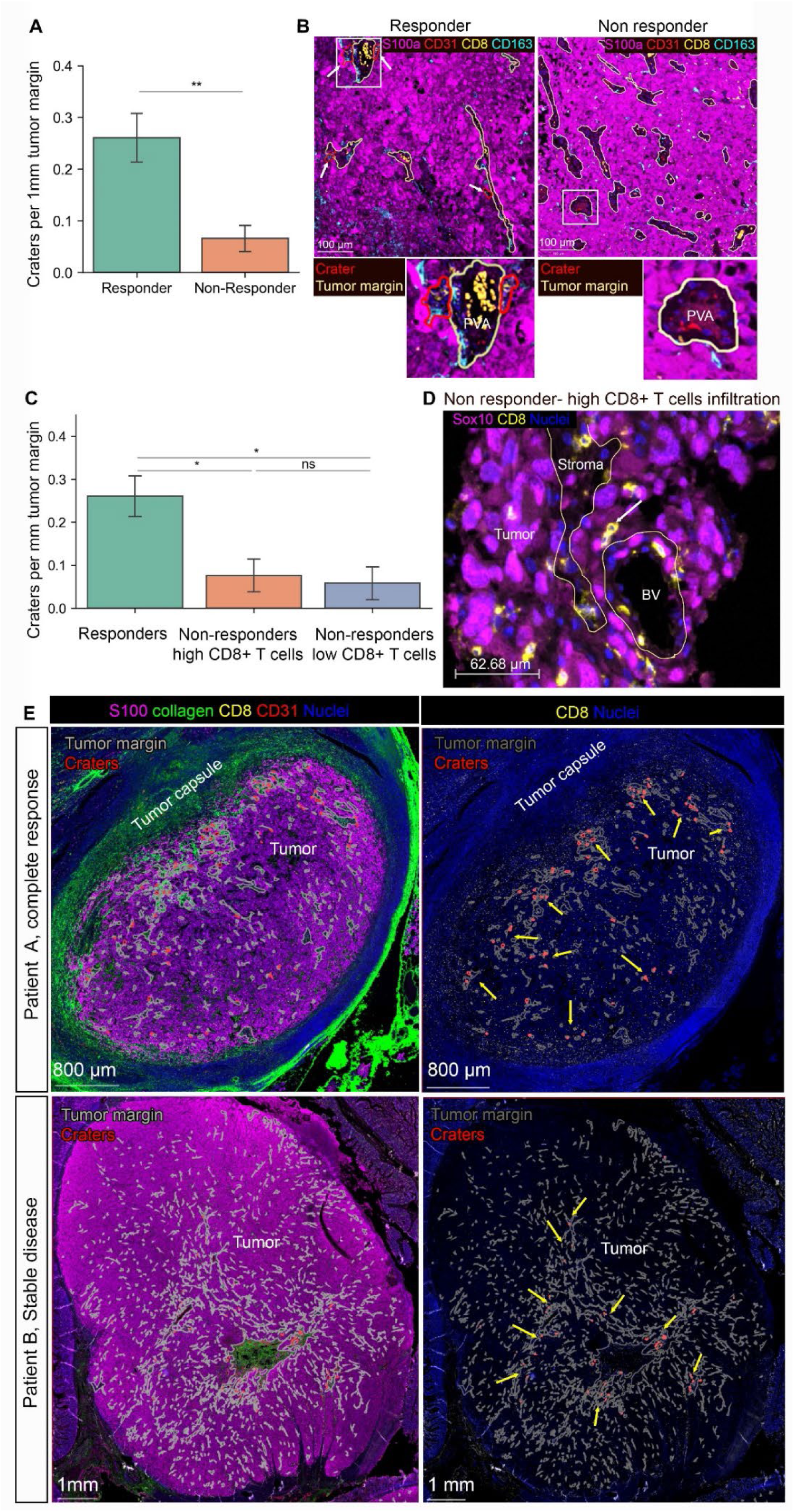
CD8^+^ T cells accumulation in craters and crater density within tumors mark successful ICB therapy response. **(A)** number of craters per 1mm tumor margin (which is the stromal-tumor interface) in samples of patients post ICB treatment (n= 3 responders -stable disease and complete remission, 7 non responders (progressive disease). Mean ±SE, TTest. *P value= 0.004). **(B)** representative images of multiplex images of a responder vs. non responder (lowly infiltrated). Peri-vascular areas are circled by yellow lines and craters are circled by red lines. Rectangle areas in the low magnification images are enlarged below each image. **(C)** Non responders were divided into highly infiltrated and lowly infiltrated samples (see Figure S13A for CD8^+^ infiltration levels, materials and methods for threshold determination). Graph showing number of craters per 1mm tumor margin in responders, non-responders with high CD8^+^ T cells infiltration and non-responders with low CD8^+^ T cells infiltration. (n= 3 responders, 3 non responders with high CD8^+^ T cells infiltration, 4 non responders with CD8^+^ T cells low infiltration. mean ±SE, TTest. P values: **= 0.0038, *=0.019). ns=0.13). **(D)** representative image of melanoma in a highly infiltrated sample taken from a non-responding patient. The tumor margin is marked with yellow lines. While many CD8^+^ T cells are evident, they infiltrate the tumor as single cells or doublets (white arrow) and very few, if any, craters are found. (**E**) Two samples of patients who responded to ICB in the adjuvant setting, showing regionality of crater formation within the tumor mass. Tumor margins (stromal/tumor border) are marked with grey lines. Craters are marked with red lines. Yellow arrows indicate foci of craters within the tumor mass. Patient A had a complete response and has foci of craters near the tumor border. Patient B had stable disease and had foci of craters at the middle of the tumor mass, near a large stromal area.

Although response to therapy has been shown to be associated with general T cell infiltration ^35^, it would be an advantage to define specific regions of T cell active tumor killing. T cells in the tumor stroma could function as cytotoxic effectors or as simply persisting in an exhausted state. The localization of CD8^+^ T cells in the immunologic niche of craters represents a more specific marker of tumor response to therapy. We found that among patients that did not respond to ICB treatment, tumors had significantly fewer craters along the tumor margin, compared tumors that have responded to treatment. Importantly, even in the presence of high CD8^+^ T cells infiltration in the tumor, craters provided more specific information for response. We separated non-responder patients into highly infiltrated and lowly infiltrated tumors and compared them to responder patients. We found that crater density remained low in non-responder patients even in the presence of high CD8^+^ T cells infiltration (Figure 7C, levels of CD8^+^ T cells infiltration are shown in S13A). Thus, the presence of craters determined response in the highly infiltrated tumors, demonstrating enhanced prediction of response by craters over general T cell infiltration. We noted that in the samples of non-responders, CD8^+^ T cells showed a more disseminated spatial distribution in the tumor parenchyma, rather than accumulation in craters (Figure 7D, S13B,C). Specifically, CD8^+^ T cells were embedded diffusely in the S100a^+^ tumor mass or entered the tumor as single cells or doublets (Figure S13B, right panel, and S13C), rather than cluster within craters (Figure S13B, left panel, S13D). This suggests that immunotherapy success is accomplished by both the creation of craters as well as the accumulation of CD8^+^ T cells within them.

To study tumor responses to ICB in other patient situations, we examined patients who were treated in the adjuvant setting. In two patients responding to treatment, we found craters in a regional distribution within the whole tumor mass. Presented in Figure 7E, patient A, who received anti-CTLA-4 treatment as a neoadjuvant therapy and achieved complete response at the time the biopsy was taken, presented craters near the edges of the tumor mass (Figure 7E, upper panel). Patient B, who received anti-PD-1 in the adjuvant setting, and achieved stable disease at the time the biopsy was taken, presented craters around a large stromal island at the middle of the tumor (Figure 7E, lower panel). Since craters are sites of anti-tumor immune response, regions abundant with craters likely represent areas within the tumor mass, where efficient immune reactivity occurs. Three patients on adjuvant therapy did not respond and had very few craters. This evaluation of craters may aid in determining treatment efficacy, as it may serve as a more direct and functionally impactful indicator of anti-tumor immune response within the tumor.

Lastly, we obtained a 3D reconstruction of human melanoma using reflectance confocal microscopy (RCM) imaging done in vivo, in patients^36^. RCM in untreated tumors exhibited crater-like structures resembling zebrafish craters, harboring lymphocyte-like cells (Figure S14A, video 6). A patient who underwent T-VEC treatment displayed enlarged crater-like areas that harbored multiple lymphocyte-like cells (Figure S14B). These results support our findings that, as in zebrafish, human melanoma exhibits craters containing CD8^+^ cells.

To learn whether craters are present in other solid tumors, we reviewed fifteen samples of non-small cell lung cancer (NSCLC) at different stages of disease. As in melanoma, we noted clusters of CD8^+^ T cells with PD-L1-expressing dendritic-shaped cells in pockets breaching into the CYTOK^+^ tumor mass in large volume tumors (Figure S15B), but not in earlier stages (Figure S15A). This indicates that the formation of craters during CD8^+^ T cell infiltration is not confined to melanoma but may rather be a general feature of T cell infiltration into a solid tumor mass.

## Discussion

Melanoma, one of the most immunogenic malignancies, often shows partial and sometimes complete immunologic regression. Recent advances in targeted immunotherapies hold promise for abrogating even metastatic disease. Study of the spatial and functional dynamics of immune cell interactions with melanoma cells within tumors has been limited by current imaging techniques^37^. Using the zebrafish, we were able to combine wide field, 3D imaging of endogenous melanoma tumors with extended time-lapse imaging in live, anesthetized, and intact animals, and record natural immune-tumor interactions for over 15 hours. We found that CD8^+^ T cells accumulate and form prolonged interactions with melanoma cells within specific structures, craters, which breach into the tumor mass from its margins. Characterization of the craters in both zebrafish and human melanoma revealed that these structures are sites that retain higher levels of antigen presenting molecules, compared to elsewhere in the tumor. Specifically, we found locally high expression of MHC class I, B2M, in melanoma cells lining craters in which CD8^+^ T cells were localized in zebrafish and human tumors. In human melanomas, the craters were also sites with an abundance of antigen presenting DCs. High levels of MHC class II and melanoma antigen MART-1 were present in DCs within craters of human tumors, consistent with their function as sites of antigen uptake and presentation by DCs. This was accompanied by the identification of CD8^+^ DCs within craters in zebrafish. Taken together, craters were identified as physical domains where the tumor is highly visible to CD8^+^ T cells, which engage it. This marked the craters as sites prone to immune attack by CD8^+^ T cells and suggested to us that once CD8^+^ T cells are activated, as during immunotherapy, craters will serve to propagate anti-tumor immune response, as antigen recognition is an essential step in CD8^+^ T cell mediated tumor killing ^38^.

Indeed, following immune stimulation, the craters serve as hotspots for an effective immune response against the tumor. The craters harbored IFN-γ^+^, activated CD8^+^ T cells and became major sites of tumor cell cytotoxic death in zebrafish. Moreover, crater size and number enlarged following CpG ODN immunostimulatory treatment. In accordance, following ICB treatment, crater density along the tumor margins was higher in patients responding to treatment, compared to non-responders, even when CD8^+^ T cells infiltrated non-responding tumors. This suggests that crater density at the tumor margin is a functionally informative pathological biomarker for an efficient anti-tumor immune response to ICB therapy. To date, efficacy of therapeutic response to immunotherapy is assessed mainly by estimating the amount of tumor necrosis and fibrosis. These features are crude and likely miss a central issue: direct evidence of ongoing therapy-enhanced immune cell-tumor cell interaction. Indeed, because durable responses to ICB may last long after cessation of therapy, recognition of the presence and quantity of specialized microdomains that inform host anti-tumor responsiveness could provide a key new tool in the predictive armamentarium of both melanoma pathologists and clinicians. Thus, crater density along the tumor-stromal margins and their regional distribution may potentially serve as an early indicator of ongoing effective anti-tumor immune responses, as well as assist in determining or modifying treatment regimens in order to further enhance the efficacy of ICB therapy.

T cells themselves appear to play a role in crater remodeling. T cell-deficient fish presented a marked reduction in crater coverage on the tumor surface. Further, crater coverage significantly increased in tumors following CpG ODN treatment, harboring high numbers of activated CD8^+^ T cells. In addition, we noted reduced numbers of craters in “cold” tumors, presenting low levels of infiltration. This indicates that immune response itself can shape tumor architecture through apoptosis that is spatially restricted to craters, and thus create focal immune regulatory sites that will affect its further propagation. We found accumulation of dead or dying cells within the enlarged craters following CpG ODN treatment and observed direct interactions between CD8^+^ T cells and apoptotic melanoma cells at the crater border in zebrafish. In addition, monocytes were found in the craters, some engulfing melanoma cells, suggesting that the dying melanoma cells were being cleared from the craters in a manner typical of apoptotic cell injury. Melanoma cell death and clearing may thereby remodel the crater structure in untreated tumors and promote gradual attrition of melanoma cells during immune-mediated regression with immunotherapy. Thus, our findings further suggest that local immune response may initiate the formation of craters. Several studies have indicated that the immune response within tumors is regional, reporting recurrent clustering of T cells with DCs and macrophages in multiple cancer types^3–5^. Such immune sites may initiate crater formation by forming a local inflammatory response that creates a focal attrition of the tumor. We noted IFN-γ expression, a marker of activation, in CD8^+^ T cells following treatment concomitant with crater enlargement. IFN-γ by itself was reported to enhance T cell cytotoxicity and motility to skin cells that express OVA in an autocrine manner in a model of graft rejection^39^. Thus, it may further contribute to crater remodeling. Moreover, IFN-γ can elevate the expression of PD-L1^40^ and MHC class I^41^, both found to be highly expressed by the DCs occupying craters in human melanoma. Therefore, local inflammation and IFN-γ secretion may have a function in the establishment of the crater’s characteristics as seen in this study.

Increased antigen recognition within craters underlies CD8^+^ T cells localization to craters and propagate tumor killing within the craters. However, it is likely that craters also harbor other factors that regulate CD8^+^ T cell arrival to craters, fate, and activity. The closed compartment created by crater architecture may facilitate a focal concentration of cytokines that affect CD8^+^ T cells within the tumor.

Moreover, crater formation is not limited to primary melanoma. In human samples, we identified craters in lung cancer, indicating that craters are a phenomenon accompanying T cell infiltration in multiple tumor types and tissue sites.

In summary, human melanomas, like those of the zebrafish, often exhibit an early, well-defined pattern of superficial (radial) growth accompanied by an underlying immune response. Over time, lesions progress to invasive, tumorigenic phases where the tumor-stromal border becomes an interface for a variable immune response by TILs. While such responses have been categorized as brisk, non-brisk, or absent, and have been linked to patients’ survival^42^, little has been known regarding their spatial organization and its significance. Craters, as described herein, are novel pathological structures representing spatially restricted and specialized hubs of the anti-tumor immune response. Improved understanding of how tumor-immune microenvironmental structures such as craters may be leveraged to affect a more robust anti-tumor immune response may ultimately facilitate immunotherapy for melanoma as well as for other tumors. Moreover, robust clinical trials should follow to assess craters as an early pathological marker for ICB treatment efficacy, which is currently not available in the clinic. This will serve to optimize clinical treatment regimens to enhance ICB success and patient well-being.

## Acknowledgments

We thank Yan Chuan and Dr. David M. Langenau for providing the Jak3 transgenic fish and thank Dr. Langenau for also kindly providing the lck promoter. We thank the FACS facility (Z. Niziolek, Jeffery L. Nelson) for their help in flow cytometry guidance, acquisition and sorting, the Bauer core facility (Claire Rearson Hartmann and Nicole Ramirez), the Harvard Center for Biological Imaging (RRID:SCR_018673) for infrastructure and their guidance. The Neurobiology Imaging facility (Mahmoud El-Rifai) for RNA Scope preparations (NIF is supported by HMS/BCH center for Neuroscience research #NS072030). We thank Drs. Melissa Gill, Salvador Gonzalez, and Christi Fox (MSK, New York) for pathological interpretation of reflective confocal microscopy images. We thank HSCRB Fish facility (Isaac Adatto, Lauren Krug, Shannon Fryer and Hannah Nations) for fish maintenance and handling. We thank Ms. Kelly Finan for creating the graphical illustration of human craters. We thank Dr. Kai W. Wucherpfennig for valuable comments and discussion, and the Zon lab members for helpful discussions and reagents. This work was supported by MRA Established Investigator Award #622785 (LIZ), NIH Fisher Program Project 5P01CA163222 (LIZ) and Cycle grant MSK project number C21883253 (LIZ). AL was supported by the Human Frontier Science Program post-doctoral fellowship LT000446/2014. GLS was supported by F31 grant CA260802. Imaging was partially supported by the 2018 Simmons award (GLS). AJN was supported by grant R00CA256497. DHK was supported by a Fred Lovejoy Resident Research and Education Award and a BCH MSO House Staff Development Award.

## Author contributions

AL conceptualized the study, wrote the manuscript, created the Tg(CD8:EGFP), Tg(lck:mCherry) transgenes and their derivate transgenic zebrafish, developed and performed long term time lapse imaging, performed experiments, analyzed data and acquired funding. GLS designed and performed experiments and wrote the manuscript, performed and analyzed single cell RNA-Seq data, implemented B2M gRNA to plasmid, prepared and analyzed samples for spatial transcriptomics assay and acquired funding. AT designed and built the long-term imaging water flow system, created image analysis algorithms and analyzed imaging data. AJN acquired CyCIF human samples and analyzed samples. AC, NB, SJ and KP collected, stained and acquired images of human samples by multiplex imaging. MM identified and provided human samples. SL, IB, QG performed and analyzed Slide-seq spatial transcriptomics. CPR identified, verified and generated a functional B2M gRNA. AS acquired and analyzed reflectance confocal microscopy images. EJ performed and analyzed single cell RNA-seq data. JVA, BR provided NLCLC samples. DSR guided imaging data acquisition and analysis. JW, HM, MES, AA, HLC, JS, WDM, EM performed experiments. SY and YZ analyzed RNA and ATAC Sequencing data, and reviewed the MS. MC, LD recruited patients and acquired RCM data, MR provided reflectance confocal microscopy data, interpreted RCM data. SRQ provided single cell RNA-seq data acquisition, MMA provided NLCLC samples. FC supervised Image Seq spatial transcriptomics. PKS, FSH, SJR and GFM provided human samples, conceptualized and designed the human data study and wrote the manuscript. LIZ conceptualized the study, supervised the study, wrote the manuscript and acquired funding.

## Methods

### Zebrafish husbandry

Zebrafish (*Danio rerio*) were bred and maintained in accordance with Animal Research Guidelines of the Institutional Animal Care and Use Committee at Harvard University (protocol #11-21), following standard protocols (at 28.5°C water temperature and a 14/10-hour light/dark cycle). Individual mating of the transgenic fish-Tg(*cd8α*:EGFP) and Tg(*lck*:mCherry) (generation described below), Tg(*flk*:EGFP)^1^, Tg(*fli1ep*:dsredex)^2^, or Tg(BRAF^V600E^/p53^null^/nacre^null^)^3^ were done to create double transgenic zebrafish lines. Jak3^P369fs^ heterozygous zebrafish (kindly provided by David Langenau) and prkdc^D3612fs^ were in-crossed, injected with tumor constructs using the MAZERATI approach (described below), and genotyped as previously described^4^. All Jak3^P369fs^ zebrafish were raised on a standalone rack with an antibiotic mix added weekly (dosage per liter fish water): 25mg Penicillin G Sodium (Sigma Aldrich, P3032), 36mg Streptomycin Sulfate (Sigma Aldrich, S6501), 0.1mg Amphothericin B (Santa Cruz Biotechnology, SC-202462B), 8mg Keflex/Cefalexin (Fish Mox Fish Flex), 65uL Prazipro/Praziquantel (Hikari Usa, AHK73254).

### Generation of *cd8a*:EGFP transgenic zebrafish

A 3878 base pair sequence upstream of the zebrafish CD8a gene, that contains open chromatin regions identified in ATAC-Seq of lck:GFP^+^ cells, was cloned from genomic DNA of zebrafish obtained from adult zebrafish of AB strain, using the following sequences: Fw: CGAGCTGTAAACACACACAA Rv: TGTTTGCTGTGAGTAACGTG. The fragment was incorporated into a pENTR 5’ TOPO TA vector and was used to drive EGFP expression using the multisite Gateway cloning technology, with the pDestTol2CG2 as a backbone plasmid, which induces EGFP expression in the heart. This facilitated selection of zebrafish in which the plasmid integrated into their genome. Single-cell embryos of casper background zebrafish^5^ were injected with the plasmid and Tol2 mRNA at 1:1 ratio. Transgenic founders were selected based on EGFP fluorescence in the heart. Fish were bred for three generations to create stable transgenic zebrafish.

### Alexa Flour 594 conjugate Ovalbumin uptake assay

Ovalbumin Alexa Flour 594 conjugate (Invitrogen) was dissolved in PBS to a 5mg/ml stock concentration. 5ug OVA-AF594 was injected intra peritoneal at a volume of 4ul to anesthetized Tg(CD8α:GFP) zebrafish weighing ∼880mg. 24 hours post injection, the kidney marrow was harvested from either OVA-AF594 injected or PBS injected zebrafish, analyzed and sorted using FACs Aria III (Becton Dickinson), as described above, to be used for inverted confocal imaging, or the zebrafish thymus was imaged by confocal in vivo imaging as described above.

### Generation of *lck*:mCherry and (*cd8α*:EGFP;*lck*:mCherry) transgenic zebrafish

Lck reporting transgenic zebrafish were produced by using the 5.5-kb sequence upstream of the *lck* start codon (kindly provided by David Langenau)^4^ to drive mCherry expression, using the multisite Gateway cloning technology ^6^. The vector was generated using the pDestTol2pA2 backbone vector. Single-cell embryos of Tg(*cd8*α:EGFP) zebrafish or of casper zebrafish were injected with the plasmid and Tol2 mRNA at 1:1 ratio, to create double transgenic or single lck:mCherry transgenic fish respectively. Transgenic founders were selected based on mCherry fluorescence in the thymus.

### Generation of melanoma tumors

Melanoma tumors were generated using two approaches. The first approach used injection of a MiniCoopR vector carrying an mitfa mini-gene and mCherry expressed under the mitfa promoter, together with Tol2mRNA at 1:1 ratio, into a single cell-stage embryo of Tg(*cd8α*:EGFP;BRAF^V600E^;p53^null^;nacre^null^) zebrafish, created by breeding of the newly generated cd8a:EGFP line described above with the establ^ished^ (BRAF^V600E^;p53^null^;nacre^null^) line, followed by genotyping to identify ger^mline^ presence of BRAF^V600E^ and the p53^null^;nacre^null^ mutation in the newly created line, as previously described^7^, and propagate the fish line as a stable line. This When use of a zebrafish line other ^than^ the Tg(*cd8α*:EGFP;BRAF^V600E^;p53^null^;nacre^null^) was desired, fluorescently-labeled, non-pigmented melanoma was generated using the MAZERATI system^8^. Single cell-stage embryos were injected with a cocktail of three vectors: a MiniCoopR v^ector^ expressing an mitfa mini-gene and BRAF^V600E^, a CRISPR MiniCoopR vector expressing an mitfa mini-gene, mitfa-driven Cas9, a gRNA targeting Tyrosinase (GGACTGGAGGACTTCTGGGG), and a gRNA targeting p53 (GGTGGGAGAGTGGATGGCTG), and a vector expressing BFP or mCherry under the mitfa promoter. We did not find a significant difference in T cell behavior and distribution between these two approaches, and our findings were corroborated in tumors generated by either approach.

### Tissue harvesting, flow cytometry and sorting

Zebrafish were euthanized in ice water. Thymus, spleen and kidney marrow or melanoma tumors were harvested under fluorescent stereo microscope Leica M165 FC. The tissues were meshed into a single cell suspension in HBSS medium (Gibco) enriched with 0.2% Fetal calf serum (FCS, Atlanta Biologicals), using a pestle. The samples were then filtered through a 40 µm filter, centrifuged at 470g for 5 minutes. The supernatant was removed, and the samples were resuspended each in 400µl HBSS+0.2% FCS. Samples were analyzed and cells were sorted into 2ml of HBSS+0.2%FCS on a FACs Aria III (Becton Dickinson) with a 100 µm nozzle at 22psi, using the FACSDiva software (Becton Dickinson).

### RNA collection and extraction

Fluorescent tagged cells sorted to purity of over 90% were used for RNA sequencing. The cells were centrifuged at 470g for 5 minutes, supernatant was removed and the cells were re suspended in 500 µl Trizol (Invitrogen) and kept in −20°C until RNA extraction. RNA was extracted using small scale RNA isolation protocol. Briefly, samples were thawed to RT and chloroform was added at 1:5 ratio. Samples were then centrifuged at 12,000g for 15 minutes at 4°C and the aqueous phase containing the RNA was collected and isopropanol was added.

Samples were kept over night at −20°C and were centrifuged at 12,000g for 10 minutes at 4°C to collect RNA. The RNA pellet was washed 3 times with 75% ethanol in nuclease-free water, re-suspended in 10 µl nuclease-free water and kept in −80°C until use.

### Ultra low input RNA sequencing (bulk RNA-seq)

CD8α:EGFP^+^ cells were sorted from thymi of 7 weeks (1.5 months old) and 12 weeks (3 months old) old CD8α:EGFP transgenic zebrafish or from melanoma from melanoma bearing Tg(*cd8α*:EGFP;BRAF^V600E^;p53^null^;nacre^null^) zebrafish. 1 ng of RNA was used from each sample. cDNA was prepared using SMART-Seq® Ultra^TM^ low Input RNA kit for sequencing (Clontech) according to the manufacturer’s instructions. Libraries were prepared using Nextera XT DNA library preparation kit (Illumina) and Nextera XT index kit (Illumina) according to the manufacturer’s instructions. Libraries prepared went through quality control analysis using an Agilent Bioanalyzer. Samples with appropriate nucleosomal laddering profiles were selected for next generation sequencing using Illumina Hiseq 2500 platform.

### RNA-seq analysis

Quality control of RNA-Seq datasets was performed by FastQC and Cutadapt to remove adaptor sequences and low-quality regions. The high-quality reads were aligned to UCSC build danRer7 (lck:GFP^+^ RNA Seq) or GRCz10 (CD8:GFP^+^ RNA Seq) of zebrafish genome using Tophat 2.0.11 without novel splicing form calls. Transcript abundance and differential expression were calculated with Cufflinks 2.2.1. FPKM values were used to normalize and quantify each transcripts; the resulting list of differential expressed genes are filtered by log fold change > (0.05) and q-value > (0.05).

### Assay for Transposase Accessible Chromatin (ATAC-seq)

50,000 lck:GFP^+^ cells were sorted from thymi of 8 months old (juvenile) adult Tg(*lck*:GFP) zebrafish at purity of above 95%. Cells were washed once with 50 μl of cold 1X PBS and spin down at 500 x g for 5 min, 4°C. After discarding supernatant, cells were lysed using 50 μl cold lysis buffer (10 mM Tris-HCl pH 7.4, 10 mM NaCl, 3 mM MgCl2, 0.1% IGEPAL CA-360) and spun down immediately at 500 x g for 10 mins, 4°C. Then the cells were precipitated and kept on ice and subsequently resuspended in 25 μl 2X TD Buffer (Illumina Nextera kit), 2.5 μl Transposase enzyme (Illumina Nextera kit, 15028252) and 22.5 μl Nuclease-free water in a total of 50 µl reaction for 1 hr at 37°C. DNA was then purified using Qiagen MinElute PCR purification kit (28004) in a final volume of 10 μl. Libraries were constructed according to Illumina protocol using the DNA treated with transposase, NEB PCR master mix, Sybr green, universal and library-specific Nextera index primers. The first round of PCR was performed under the following conditions: 72°C, 5 min; 98°C, 30 sec; [98°C, 10 sec; 63°C, 30 sec; 72°C, 1 min] X 5 cycles; hold at 4°C. Reactions were kept on ice and using a 5 µl reaction aliquot, the appropriate number of additional cycles required for further amplification was determined in a side qPCR reaction: 98°C, 30 sec; [98°C, 10 sec; 63°C, 30 sec; 72°C, 1 min] X 20 cycles; hold at 4°C. Upon determining the additional number of PCR cycles required further for each sample, library amplification was conducted using the following conditions: 98°C, 30 sec; [98°C,10 sec; 63°C, 30 sec; 72°C, 1 min] X appropriate number of cycles; hold at 4°C. Libraries prepared went through quality control analysis using an Agilent Bioanalyzer. Samples with appropriate nucleosomal laddering profiles were selected for next generation sequencing using Illumina Hiseq 2500 platform.

### ATAC-seq analysis

All zebrafish ATAC-Seq datasets were aligned to build version Zv9/danRer7of the zebrafish genome using Bowtie2 (version 2.2.1) ^9^ with the following parameters: --end-to-end, -N0, -L20. We used the MACS2 version 2.1.0 ^10^ peak finding algorithm to identify regions of ATAC-Seq peaks, with the following parameter --nomodel --shift -100 --extsize 200. A q-value threshold of enrichment of 0.05 was used for all datasets.

### Imaging of still images

Gross fluorescent images were taken using a fluorescent stereo microscope Leica M165 FC equipped with a 20 megapixel color CMOS camera DMC5400, and using Leica LAS X software. Confocal Images of sorted CD8α:GFP^+^ lymphoid T cells and myeloid DCs were obtained on an LSM 880 microscope (ZEISS, Thornwood, NY). GFP was excited sequentially with a 488 nm laser line. Fluorescence was captured with a 63x 1.4 NA oil immersion objective. Laser excitation light passing through the sample was captured by a 0.55 NA condenser equipped with DIC optics and imaged onto a PMT detector to produce an overlaid DIC image.

To image zebrafish melanoma for still live imaging, the zebrafish were euthanized in 4g/L Tricaine, fixated on 35×10 mm tissue culture dish (Falcon), immersed with PBS and put directly under the scope without further manipulation. Confocal in vivo imaging of melanoma was performed on an Axio Examiner upright microscope (ZEISS, Thornwood, NY) equipped with an Insight DS+ tunable laser (Spectra-Physics). EGFP and mCherry were excited simultaneously with 930 nm light. Fluorescence was captured with a 10x 0.5 NA W Plan-Apochromat water dipping objective and imaged onto non-descanned GaAsP detectors. Image stacks were taken with a 2 mm spacing.

Second harmonic generation (SHG) Imaging to detect collagen was performed on an upright AxioExaminer microscope with LSM 980 scanhead and a two-channel non-descanned BiG.2 detector (Carl Zeiss Microscopy, Whiteplains, NY). SHG signal was generated with an InSight X3+ IR laser tuned to 930 nm. The back-scattered signal was collected by a 10x 0.5 NA W Plan-Apochromat objective and imaged through a 460-500 nm bandpass filter.

Images were processed using the Imaris software (Oxford Instruments), followed by minor brightness and contrast corrections applied to the entire image, to allow clear visualization, if needed, using photoshop software (Adobe).

### Time-lapse live imaging

To maintain the zebrafish alive for 15 hours, a water flow system was built based on Xu et al. ^11^, with modifications as follows. 0.75x Tricaine (1g/8L system salted water+E3+Tris pH 9.0) was pumped using Ismatec REGLO Digital 4-channel 12 Roller variable speed pump (Cole Parmer, USA) in a constant flow rate of 6 ml/minute into the mouth of intubated zebrafish placed on custom-made stainless-steel plate with an orifice to allow suction of excess of water during the imaging period. The zebrafish was anesthetized and stabled using 1% agarose dissolved in salted water before intubation. Throughout the imaging time, excess of water was suctioned out using a KNF N86 KTP vacuum pump (Cole parmer, USA). Water temperature was kept constant at 28°C using SH-27B in-line heater and controlled by TC-324 heater controller (Harvard Apparatus, USA). The pressure at the output was measured using ProSense digital pressure sensor (AutomationDirect, USA), and a feedback loop to the peristaltic pump using Arduino Uno microprocessor was used as a safety mechanism in case of clogging. Time-lapse live imaging was obtained using ZEISS LSM 980 scanhead coupled to an Axio Examiner upright microscope (both, ZEISS,Thornwood, NY) and equipped with an Insight DS+ tunable laser (Spectra-Physics) as described above. 3D Images were acquired every 6-8 minutes for 20 hours with stacks taken with a 2 µm spacing.

### Analysis and quantification of 3D images

Image processing was performed using Arivis Vision4D 3.4-3.6. For the segmentation of live 3D images two dedicated pipelines were used, one for the segmentation of the tumor and craters, and another for T-cells. When treatment vs. control, or gene KO vs, wild type samples were quantified, the analysis was done blind to the condition of the analyzed sample, by a different person (AT) than the one collecting the data (AL or GLS). For the tumor and craters, bright mCherry cells were enhanced using the “Normalization” function and segmented using “Blob finder” in order to exclude them from further analysis. For tumor segmentation Gaussian noise reduction was used followed by “Otsu” thresholding method. For the segmentation of craters a black top-hat transform was performed on the tumor segment, in short, morphological closing with a 40 voxels square kernel, followed by subtraction of the tumor segment from the morphological closing result, an additional step of morphological opening with a 2 voxels kernel was performed to clean up residual segments arising from the mismatch between the tumor morphology and the square kernel. For T-cells segmentation Gaussian noise reduction followed by “Blob finder” was performed on the GFP channel. All resulting segmentation were visually inspected and manually corrected if necessary. Resulting dimensions, distance calculations and an image of the segmented tumor and craters were exported to MATLAB R2020 for calculations of tumor coverage and T-cells density, calculated for CD8^+^ cells in direct contact with segmented crater area or tumor area.

### FACS sorting of T cells for scRNA-seq

Melanoma tumors from two Tg(*cd8*α:EGFP;*lck*:mCh) fish injected with tumor constructs using the MAZERATI approach^8^ (described above) and normal skin from 1 Tg(*cd8α*:EGFP;*lck*:mCh) fish were dissected and processed as previously described. Lysis buffer was prepared containing per reaction: 125U RNase inhibitor (Takara Bio, 2313A), 2.5mM dNTP (NEB, N0447S), 2.5 µM oligo-dT primer (IDT, 5′AAGCAGTGGTATCAACGCAGAGTACT30VN-3′), .05% Triton X-100 (Sigma-Aldrich, 93443), and nuclease-free water (Ambion, 9937) to 4 µl. 96-well PCR plates (Bio-Rad, HSP9601) were loaded with 4 µl lysis buffer per well, sealed with Aluminum Foil-96 One Tab seals (VWR, 60941-126), and placed on ice. Cells were stained with Sytox Red (Thermo Scientific, S34859) immediately before FACS-sorting individual live cells in the 96-well plates on a FACs Aria III (Becton Dickinson), as described above. 6 plates and 2 plates of cd8α:EGFP^+^l/ck:mCh^+^ and cd8α:EGFP^-^/lck:mCh^+^ cells were sorted from tumor and normal skin, respectively, for totals of 576 cells from tumor and 192 cells from normal skin. Plates were spun down and stored at −80°C immediately after sorting.

### Smart-seq library generation

cDNA synthesis was performed using the Smart-seq2 protocol^12,13^. In brief, 96-well plates containing single-cell lysates were thawed on ice followed by first-strand synthesis. 6 μl of reaction mix (16.7 U μl^−1^ SMARTScribe Reverse Transcriptase (Takara Bio, 639538), 1.67 U μl^−1^ Recombinant RNase Inhibitor (Takara Bio, 2313B), 1.67X First-Strand Buffer (Takara Bio, 639538), 1.67 μM TSO (Exiqon, 5′-AAGCAGTGGTATCAACGCAGAGTGAATrGrGrG-3′), 8.33 mM dithiothreitol (Bioworld, 40420001-1), 1.67 M Betaine (Sigma, B0300-5VL) and 10 mM MgCl_2_ (Sigma, M1028-10X1ML)) was added to each well. Reverse transcription was carried out by incubating wells on a thermal-cycler (Biorad) at 42°C for 90 min, and stopped by heating at 70°C for 5 min. Subsequently, 15 μl of PCR mix (1.67X KAPA HiFi HotStart ReadyMix (Kapa Biosystems, KK2602), 0.17 μM IS PCR primer (IDT, 5′-AAGCAGTGGTAT CAACGCAGAGT-3′), and 0.038 U μl^−1^ Lambda Exonuclease (NEB, M0262L) was added to each well, and second-strand synthesis was performed using the following program: 1) 37°C for 30 min, 2) 95°C for 3 min, 3) n cycles of 98°C for 20 s, 67°C for 15 s and 72°C for 4 min, and 4) 72°C for 5 min, with n=23 for plate N1 and n=26 for plates 111-11++, 107-24, 111-11 Mch, and 111-11++. An AMPure XP bead (Beckman Coulter, A63882) clean-up was performed according to manufacturer instructions, with bead:sample ratio .7X. The amplified product was diluted with a ratio of 1 part cDNA to 20 parts EB (Qiagen, 1014609), and concentrations were measured with a dye-fluorescence assay (Quant-iT dsDNA High Sensitivity kit; Thermo Fisher, Q33120) on a SpectraMax i3x microplate reader (Molecular Devices). cDNA was diluted in EB to final per-well concentration of approximately 0.25 ng/µl. Illumina sequencing libraries were prepared using 0.4 µl of cDNA from each sample well, as described previously^14^. Libraries were sequenced on the NovaSeq 6000 Sequencing System (Illumina) using 2 × 100-bp paired-end reads and 2 × 12-bp index reads with either a 200- or 300-cycle kit (Illumina, 20012861 or 20012860).

### scRNA-seq data processing

Quality control of smart-Seq datasets was performed by FastQC (http://www.bioinformatics.babraham.ac.uk/projects/fastqc/) and Cutadapt^15^ to remove adaptor sequences and low-quality regions. The high-quality reads were aligned to a Zon lab custom genome^16^ using STAR 2.7.0 Spliced Transcripts Alignment tool^17^.

Analysis was performed using Seurat v4.0.2 and R v4.0.4. Cells with fewer than 500 or greater than 3000 features and cells with greater than 10% mitochondrial reads were excluded from analysis. Cells were split into Normal and Tumor samples and batch corrected using Seurat scRNA-seq integration^18^ with 2000 integration features. Principal component analysis, clustering, and UMAP dimensionality reduction was performed using Seurat with default parameters and selection of 15 dimensions, based on plotting the percentage of variance explained by each principal component, and a resolution of 0.8. Differentially expressed gene markers were identified using the Seurat function FindAllMarkers with a minimum percent expressing cells of 0.25 and the default log-fold change threshold of 0.25. Genes included in expression plots not identified as cluster-specific markers: bcl11ba, TCRa.C, TCRb.C1, trdc, tcrg, mmp9, lyz, cd8β, and cd8α.

### Tissue preparation and FACS for vascular cells scRNA-seq

We selected an adult Tg(fli1ep:dsRedex, flk1:EGFP) zebrafish bearing an mitfa:BFP^+^ tumor generated by the MAZERATI system (injection with 1) a MiniCoopR vector expressing an mitfa mini-gene and BRAFV600E, 2) a CRISPR MiniCoopR vector expressing an mitfa mini-gene, mitfa-driven Cas9, a gRNA targeting Tyrosinase (GGACTGGAGGACTTCTGGGG), and 3) a gRNA targeting p53 (GGTGGGAGAGTGGATGGCTG), and a vector expressing BFP under the mitfa promoter). The fish bore a protruding tumor with tumor-adjacent skin that presented as normal skin (lacking BFP^+^ cells or dark pigmented melanocytes). The fish was euthanized by rapid cooling in ice water. The tumor and healthy skin were independently harvested and transferred to HBSS medium (Gibco). The tissue was dissociated using a size 11 surgical blade, then incubated for 30 minutes in 0.15mg/mL Liberase (Sigma cat. no. 05401119001) in HBSS while shaking at 150-200rpm at 37C. The solution was resuspended by pipetting halfway through the 30 minute incubation. FBS was added to 10% and cells were passed through a 40um filter (additional 500uL HBSS+2%FBS was passed through to rinse filter) and pelleted by centrifugation at 470g for 5 minutes before resuspension in HBSS+2% FBS. Cells were stained with Sytox Red (Thermo Scientific, S34859) as a live/dead stain (Abcam) and sorted on a FACs Aria III (Becton Dickinson) using the FACSDiva software (Becton Dickinson). Gates were drawn using negative and single-fluorescent reporter control zebrafish. FACS index data was recorded during sorting. Fli1a:dsRed^+^;Flk1:GFP^+^ and Fli1a:dsRed^+^;Flk1:GFP-cells were sorted from both tumor and normal skin directly into 384-well capture plates (Single Cell Discoveries) and immediately spun down 1000xg for 1 min @4C before being put directly on dry ice and promptly stored at −80°C.

### Sequencing and Analysis for vascular cells

Single-cell RNA sequencing was performed by Single Cell Discoveries and sequenced by CEL-Seq2 as previously described^19^. The data was mapped by Single Cell Discoveries to Zv11 with the addition of 5 sequences: BRAFv600e, TagBFP, dsRed, mCherry, and EGFP. Analysis was performed using Seurat v4.1.0 and R v4.0.4. Cells with fewer than 50 or greater than 2000 features, fewer than 100 RNA count, greater than 10% mitochondrial reads, or greater than 70% ERCC spike-in reads were excluded from analysis. Cells were split by the plate in which they were sorted and batch corrected using Seurat scRNA-seq integration SCTransform^18^ using the default 3000 variable features. Principal component analysis, clustering, and UMAP dimensionality reduction was performed using Seurat with default parameters and selection of 30 dimensions, based on plotting the percentage of variance explained by each principal component, and a resolution of 1.0. Differentially expressed gene markers were identified using the Seurat function FindAllMarkers with a minimum percent expressing cells of 0.25 and the default log-fold change threshold of 0.25. Genes included in DotPlot expression plot are statistically significantly enriched in the specified cluster[s] (adjusted p-value <1e-9) excepting: krt91 (cluster 0 had no statistically significant marker genes and appeared to represent a mixed population), sox7, and fli1a.

### In situ transcriptome processing via Slide-seq

For Slide-seq assay, whole zebrafish bearing melanoma tumors were frozen in OCT on dry ice and placed at −80°C. Slide-seq arrays were prepared and spatial bead barcodes sequenced following Slide-seqV2^20^ protocol using arrays created with custom synthesized barcoded beads (5’-TTT_PC_GCCGGTAATACGACTCACTATAGGGCTACACGACGCTCTTCCGATCTJJJJJJJ JTCTTCAGCGTTCCCGAGAJJJJJJNNNNNNNVVT30-3’) with a photocleavable linker (PC), a bead barcode sequence (J, 14 bp), a UMI sequence (NNNNNNNVV, 9 bp), and a poly dT tail. OCT-embedded frozen tissue samples were warmed to −20°C in a cryostat (Leica CM3050S) and serially sectioned at a 10 μm thickness (2-3 Slide-seq array replicates per sample), with consecutive sections used for hematoxylin and eosin staining and immunofluorescence staining. Each tissue section was affixed to an array and moved into a 1.5 mL Eppendorf tube for downstream processing. The sample library was prepared as previously described^20^. Libraries were sequenced using the following read structure on a NovaSeq (S2; Illumina): Read1: 42 bp; Read2: 41 bp; Index1: 8 bp, and sequences were processed as previously described^20^ using the pipeline available at https://github.com/MacoskoLab/slideseq-tools. For crater vs non crater analysis, RCTD was applied as described^21^ to identify cell types. We marked crater and non-crater areas (shown in Fig. 2A, circles) on the expression map and applied C-SIDE to perform differential expression analysis as described^22^.

### B2M CRISPR mutagenesis

Target selection for CRISPR/Cas9-mediated mutagenesis was performed using CHOPCHOP^23^. The selected sgRNA for *b2m*, TAAATCCAAACCGGGCAGCG having GC-content 55%, self-complementarity=0,MM0= 0, MM1=0, MM2=0, MM3=0 and predicted efficiency of ∼ 69% of editing. The sgRNA templates were generated using the protocol as previously described^24^. mMESSAGE mMACHINETM SP6 Transcription Kit (Invitrogen, AM1340). The gRNAs were validated using T7 endonuclease I assay in 72 hpf embryos. Primers used for detection of deletion of *b2m* are: Forward: 5’-CATCTTCGCAATACCTTAGGCT-3’ and reverse: 5’-TCAGCTAAGGTAAGTGAACAGGCCTTAATTTGGAC-3’. The selected guides were integrated into a MAZERATI vector with Cas9 expressed under the mitfa promoter allowing tissue specific B2M CRISPR in melanoma^8^. Tumor grown in B2M CRISPR vector or Tyrosinase CRISPR vector as control were imaged using upright confocal. Following imaging, the tumors were resected and evaluated for editing efficiency by deep sequencing of PCR amplicons. In brief, genomic DNA from zebrafish tissue was extracted with DNeasy blood and tissue kit (Qiagen, 69506). The CRISPR loci of the targeted genes were amplified using the primers above and barcoded with the Illumina NGS adaptor. Amplification was performed using Phusion® High-Fidelity PCR Master Mix with HF Buffer (NEB, M0530), with the following conditions: 98°C 3 minutes, [98°C 10 seconds, T annealing 10 seconds, 72°C 10 seconds] x 35 cycles, 72°C 5 minutes. T annealing was 63°C]. Then, PCR amplicons were purified on PCR purification kit columns prior to sequencing. For the analysis the sequencing reads were first trimmed for quality and aligned to the GRCz11/danRer11 assembly using Bowtie2 (B. Langmead, 2012) with the --very-sensitive setting. Mutations were quantified by the R software CrispRVariants-version 1.20 (H. Lindsay,2016) using a minimum read count of 20. The R version in this analysis was 4.1.0 (2021-05-18).

### CpG ODN treatment

CpG ODN 2007 (Invivogen, USA) was dissolved in sterile water to stock concentration of 2µg/µl. CpG ODN was further diluted using PBS to 1µg/µl. For control, sterile water were diluted in PBS at the same ratio as CpG ODN. Either control or CpG ODN solution was injected directly into the tumor at a volume of 1µl using 10 µl 1701Hamilton syringe model 1701 (33s/0.6’/2). CpG ODN or control were injected daily and tumors were analyzed 24 hours post injection.

### TGF-β inhibition treatment

SB431542 (Sigma Alderich, USA) was used to inhibit TGF-β. SB431542 was dissolved in DMSO to 10mM. SB431542 was further dissolved in fish water to a concentration of 10µM. To achieve TGF-β inhibition, zebrafish containing tumors were immersed in fish water contained 10 µM SB431542 and analyzed 24 hours after treatment. As control, zebrafish were immersed in fish water containing DMSO 0.1%.

### RNA Scope

Zebrafish tumors were excised and were immediately frozen in OCT in 2-methylbutane over liquid nitrogen. Samples of 10µm were cut using cryostat (Leica) and placed on superfrost histological slides (Fisher scientific, USA) and stored in −80°C until use. RNA Scope was performed using RNAScope^TM^ Fluorescent multiplex assay (ACDBio, USA) according to manufacturer’s instructions for fresh frozen samples. RNA probes for mCherry (RNAscope-Probe-mCherry, C2 channel, stained with Atto647N), zebrafish CD8α (RNAscope® Probe-Dr-cd8a-C3 channel, stained with Atto550) were used to detect the melanoma and CD8^+^ cells. In addition, costume probes designed for the zebrafish genes ifng1, tox and prf1.1 were used to detect activation markers at the C1 channel, stained with AF488. Slides were imaged using an inverted AxioObserver with LSM 900 scanhead microscope, and equipped with 405, 488, 561 and 640 nm diode lasers. Images were prepared using Imaris software (Oxford Instruments).

### Whole mount TUNEL assay

TUNEL assay was performed using Click-iT™ Plus TUNEL Assay Kits for In Situ Apoptosis Detection (Fisher Scientific, USA) on whole tumors. Tumors were excised with wide margins from the zebrafish and incubated in 4% PFA in PBS (Boston BioProducts, USA) for 15 minutes. Samples were washed with BSA 3% in PBS twice and incubated in proteinase K solution for 20 minutes at room temperature. Washed the samples with PBS and immersed in TdT reaction buffer for 10 minutes in 37°C, then removed the reaction buffer and added the TdT reaction mix, prepared according to protocol for 500 µl to cover the samples. Incubated for 1.5 hours in room temperature. Washed the slides with 3% BSA and 0.1% Triton-X-100 in PBS for 5 minutes, then washed with PBS. To stain, added Click-iT Plus TUNEL reaction cocktail prepared according to manufacturer’s instructions for 500 µl solution and incubated for 60 minutes in 37°C incubator. Washed with 3% BSA in PBS for 5 minutes and stored at 4°C. Samples were imaged the next day in PBS using an Axio Examiner upright microscope (ZEISS, Thornwood, NY) equipped with an Insight DS+ tunable laser (Spectra-Physics). Fluorescence was captured with a 10x 0.5 NA water dipping objective and imaged onto non-descanned GaAsP detectors. To assure the signal report EdU uptake, tumor of zebrafish underwent the same staining procedure, except that PBS was added to the TdT reaction mixture instead of the TdT enzyme. Only low background signal was detected in this tumor.

### Quantitative analysis of CyCIF processed human melanoma samples

CyCIF data was procured from database collected at Nirmal et al.^25^, made available for us by AJN and PKS. The data was analyzed to identify craters in human melanoma and immune cell characteristics related to craters. Preprocessing and segmentation of the CYCIF images were performed as follows. Prior to segmentation, the membrane labeled channels (CD4, CD8a, CD163, CD3d, CD11c, MCAM, ICOS, HLAA, CD31, MART1, CD16) and the cytoplasmic label S100A were each normalized to their 99th percentile value and summed. The Nucleus channel was normalized separately. Segmentation was then performed using Cellpose [1] with the TissueNet model, allowing identification and segmentation of individual cells (Fig. S8A). For gene expression data, background subtraction was performed on each channel using a rolling ball algorithm with a radius of 50 pixels. Mean fluorescence intensity per cell was then extracted from each channel for further analysis.

Clustering and analysis of the single cell data was performed using the Scanpy [2] python package. For cell type identification, the following marker genes were utilized: SOX10, CD163, CD3d, CD8a, FOXP3, CD11c, CD4, CD68, CD31, Granz.B, and MART1. The log transformed gene expression data was scaled using standard scaling prior to dimensionality reduction. Embedding of the data was performed using UMAP^26^ and clustering was done with Leiden clustering^27^. The clustered data was then manually inspected and corrected. Clusters were then classified to specific cell types (Fig. S8B).

Crater regions were manually labeled based on the S100A, CD8a, CD31 and aSMA channels. Segmentation of the tumor mass was performed by applying Otsu thresholding to the S100A channel, followed by morphological closing to fill small holes within the tumor region, and morphological opening to remove small, disconnected tumor fragments. The tumor margin region, enclosing the S100a^+^ cell mass, was segmented by dilating the tumor segmentation and subtracting the original tumor area, defining the boundary area (Fig. S8C).

To calculate cell densities, cells overlapping the border region but not craters were classified as border cells. Tumor cells were defined as overlapping the tumor segmentation without overlapping craters or the border. This enabled quantitative analysis of cell densities across different spatial regions.

### Human sample acquisition and multiplexed immunofluorescence

Tumor samples were collected from cancer patients receiving clinical immunotherapies at Dana-Farber Cancer Institute. All individuals gave written informed consent to utilize their leftover tumor samples in accordance with the Institutional Review Board (IRB) approved protocol 17-000. Staining was performed overnight on a BOND RX fully automated stainer (Leica Biosystems). Tissue sections of 5-μm thick FFPE were baked for 3 hours at 60°C before loading into the BOND RX. Slides were deparaffinized (BOND DeWax Solution, Leica Biosystems, Cat. AR9590) and rehydrated with series of graded ethanol to deionized water. Antigen retrieval was performed in BOND Epitope Retrieval Solution 1 (pH 6) (Leica Biosystems, Cat. AR9961, AR9640) at 95°C. Deparaffinization, rehydration and antigen retrieval were all pre-programmed and executed by the BOND RX. Next, slides were serially stained with primary antibodies. Each primary antibody was incubated for 30 minutes. Subsequently, anti-mouse plus anti-rabbit Opal Polymer Horseradish Peroxidase (Opal Polymer HRP Ms + Rb, Akoya Biosciences, Cat. ARH1001EA) was applied as a secondary label for a 10 minute incubation. The corresponding Opal Fluorophore Reagent (Akoya) was applied for 10 minutes. Slides were incubated in Spectral DAPI solution (Akoya) for 10 minutes, air dried, and mounted with Prolong Diamond Anti-fade mounting medium (Life Technologies, Cat. P36965) then stored in a light-proof box at 4 °C prior to imaging. Image acquisition was performed using the Vectra Polaris multispectral imaging platform (Vectra Polaris, Akoya Biosciences). The target antigens, antibody clones, dilutions, and antigen retrieval conditions are listed in Supplementary Table 2. Images were either assessed pathologically or underwent quantitative analysis.

### Quantitative analysis of multiplexed immunofluorescence processed human melanoma samples post treatment

Identification of perivascular border was performed in a semi-automated manner using Arivis4D and QuPath. In short, The S100a channel was filtered using a median filter, followed by thresholding and segmentation of the tumor mass. The segmented area was subject to morphological closing, followed by subtraction of the non-closed segment from the closed one, resulting in segmentation holes in the tumor mass. These holes were then filtered based on size and CD31 expression to identify perivascular area. Segmentation maps were then transferred to QuPath for manual correction and manual crater identification. Perivascular boundary length was calculated using QuPath and crater density within the tumor was calculated as crater number per linear 1mm length of tumor margin. Areas of massive fibrosis were omitted from the analysis due to the inability to determine a clear stromal-tumor border.

Intra-tumoral CD8^+^ T cells density was measured and calculated at the pathology department of Brigham and Women’s Hospital (BWH), as part of sample processing for pathological evaluation. The values reflect the average of CD8^+^ T cells number per mm2 in 6 fields of view per sample and their corresponding standard error.

A cohort of 114 melanoma cases quantified for intratumoral CD8^+^ density had a median density of 258 cells/mm2 (BWH, Scott Rodig et al. unpublished), which was used as the threshold to determine high vs. low levels of inflammation.

### In vivo Reflectance confocal microscopy (RCM) imaging in human patients

In vivo patient imaging was conducted on melanoma lesions in 2 patients for clinical diagnosis or treatment monitoring after receiving T-Vec injection. Written informed consent was obtained where necessary. Imaging was performed using an RCM device (tissue-coupled VivaScope 1500 or handheld VivaScope 3000, Caliber I.D., Rochester, NY). RCM images were acquired at multiple locations within the lesion for comprehensive lesion evaluation. Between 3–4 mosaics were acquired at epidermis, dermal–epidermal junction and dermis, followed by stacks and videos. Images were read and interpreted in real-time at the bedside to select the targeted biopsy site(s) by 2 investigators (M.C. and A.S.) with over 6 years of RCM reading experience. Images were evaluated for previously described criteria for nodular melanoma^28^. Craters were defined as hypo-reflective structures within or surrounding cerebriform nests (that represent melanoma tumor nests) with absence of any blood flow. Small, low to medium reflective mononuclear cells inside and at tumor periphery were termed as lymphocyte-like cells based on previous sudies^29^.

### Statistical analysis

Statistical tests were performed using Python SciPy package v1.9.3. All fields of view from the same fish were pooled together and each fish is considered as an independent repeat. We used two sided unpaired t-test for comparison between groups with normal distribution or small sample size, and two sided Mann-Whitney U-test for groups with non-normal distribution. P < 0.05 was considered significant; *P < 0.05, **P < 0.01, ***P < 0.001.

