## Supplemental figures and tables for "Craters on the melanoma surface facilitate tumor-immune interactions and demonstrate pathologic response to checkpoint blockade in humans"

**Supplemental information:**

Figures S1-15.

Tables S1-2

Videos S1-6:

Video S1. **A $CD8^+$ T cell interacts with a melanoma cell in a crater in untreated melanoma tumor.** Related to Figure 1B.

Video S2. **$CD8^+$ T cells linger in a crater in untreated melanoma tumor.** Related to Figure S4A.

Video S3. **$CD8^+$ T cells interact with melanoma cell in a manner characteristic to tumor killing in vivo.** Related to Figure 1E.

Video S4. **$CD8^+$ T cells interact with melanoma cells in enlarged craters following CpG ODN treatment.** Related to Figure 3.

Video S5. **mCherry fragment uptake by $CD8^+$ dendritic cell following TGF-β inhibition.** Related to Figure S8D.

Video S6. **3D projection of human melanoma tumor nest, containing a crater-like structure, in a live patient.** Related to Figure S14

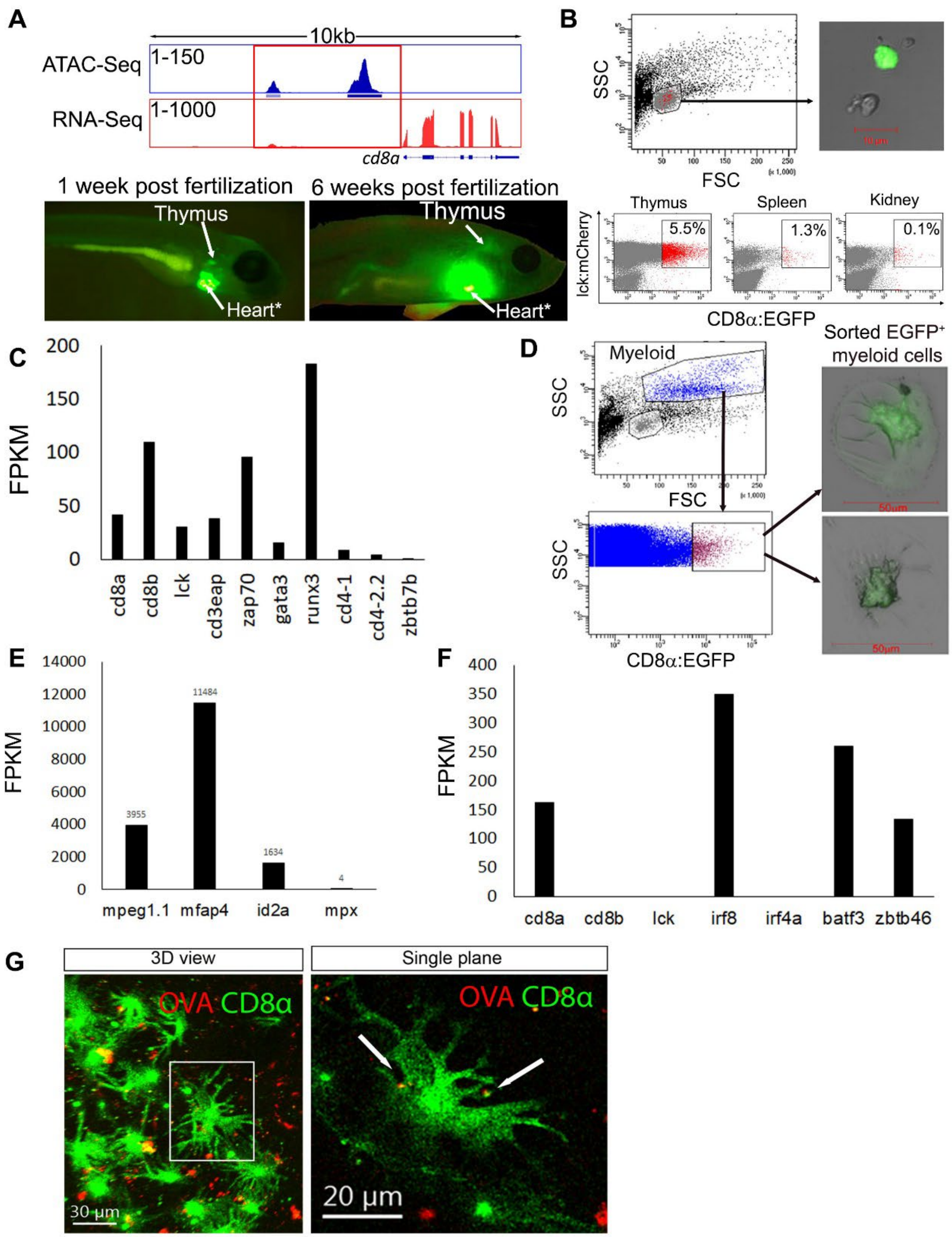


Figure S1. **Generation of cd8α:EGFP transgenic zebrafish line that reports CD8α$^+$ T and dendritic cells. (A)** Upper: the *cd8a* gene area displaying the ATAC-seq and bulk RNA-seq analyses of sorted lck$^+$ lymphocytes from zebrafish thymi. The rectangle marks the cis-regulatory element used as *cd8a* promoter to drive cell-specific transgene expression, containing two open chromatin peaks. Lower: fluorescent images of zebrafish at 1- and 6-weeks post fertilization. Arrow indicating the EGFP$^+$ zebrafish thymi. *EGFP expression in the heart is irrelevant for *cd8a* expression. It results from the backbone plasmid used to select successful transgenes. (**B**) Flow cytometry analysis of thymus derived cells from Tg(cd8α:EGFP;lck:mCherry) zebrafish. Upper plot: FSCC plot of zebrafish thymus cells. In red, EGFP$^+$ gated population appears at the lymphoid gate. Lower plots: Lymphoid gated flow cytometry plots of thymus, spleen and kidney cells, showing CD8α:EGFP$^+$/lck$^+$ cells in the major zebrafish lymphoid organs. Upper right image: EGFP$^+$ lymphocytes sorted onto slides showing a bright EGFP$^+$ cell of ~10µm in diameter. (**C**) Bulk RNA-seq of EGFP$^+$ thymus-derived sorted cells from 6 weeks old (juvenile) cd8α:EGFP transgenic zebrafish, reveals expression of genes characteristic of CD8$^+$ T cells. (**D**) Flow cytometry plots of thymus-derived cells showing cd8α:EGFP$^+$ cells in the myeloid gate. When sorted onto slides this population was comprised of large, 70µm, EGFP$^+$ cells. (**E**) Bulk RNA-Seq of EGFP$^+$ myeloid cells sorted from melanoma tumors showing a gene profile characteristic to monocytes and specifically to (**F**) CD8α dendritic cells. (**G**) In-vivo imaging of the thymic region of Tg(cd8α:EGFP;lck:mCherry) zebrafish injected I.P. with ovalbumin-AlexaFlour 594 shows cd8α:EGFP dendritic cells (3D view, left) with ovalbumin presented on their dendrites (single plane, right).

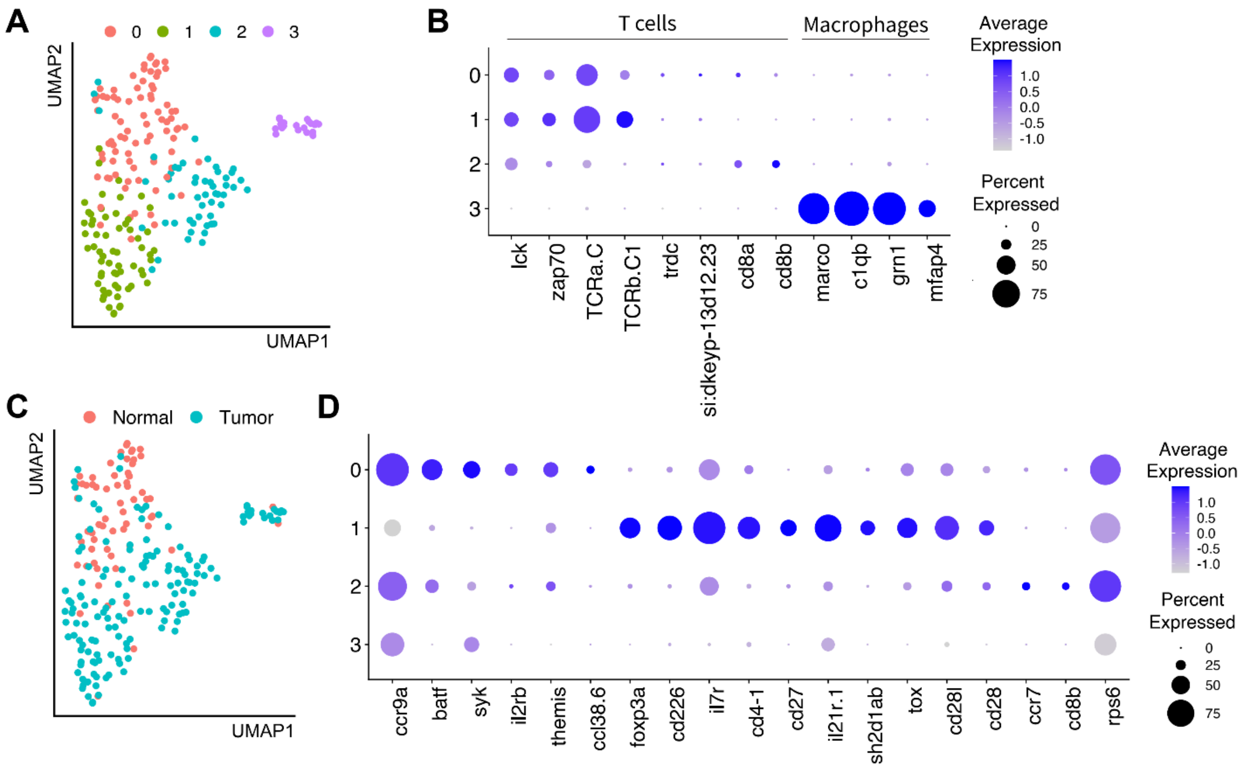


Figure S2. **Immunological landscape of melanoma tumor in zebrafish.** (**A**) Single-cell RNA-Seq by Smartseq2 of sorted T cells from zebrafish melanoma and normal skin exhibit 4 clusters. (**B**) Expression of select genes classifying T cell clusters and a cluster of macrophages. (**C**) Tissue origin labeling reveals one cluster of predominantly normal skin-derived T cells and two clusters of primarily tumor-derived T cells. (**D**) Differential gene expression analysis identifies a heterogeneous population of T cells comprising the normal skin, with tumor-derived clusters exhibiting $CD8^{+}$, $CD4^{+}$ and T regulatory cells.

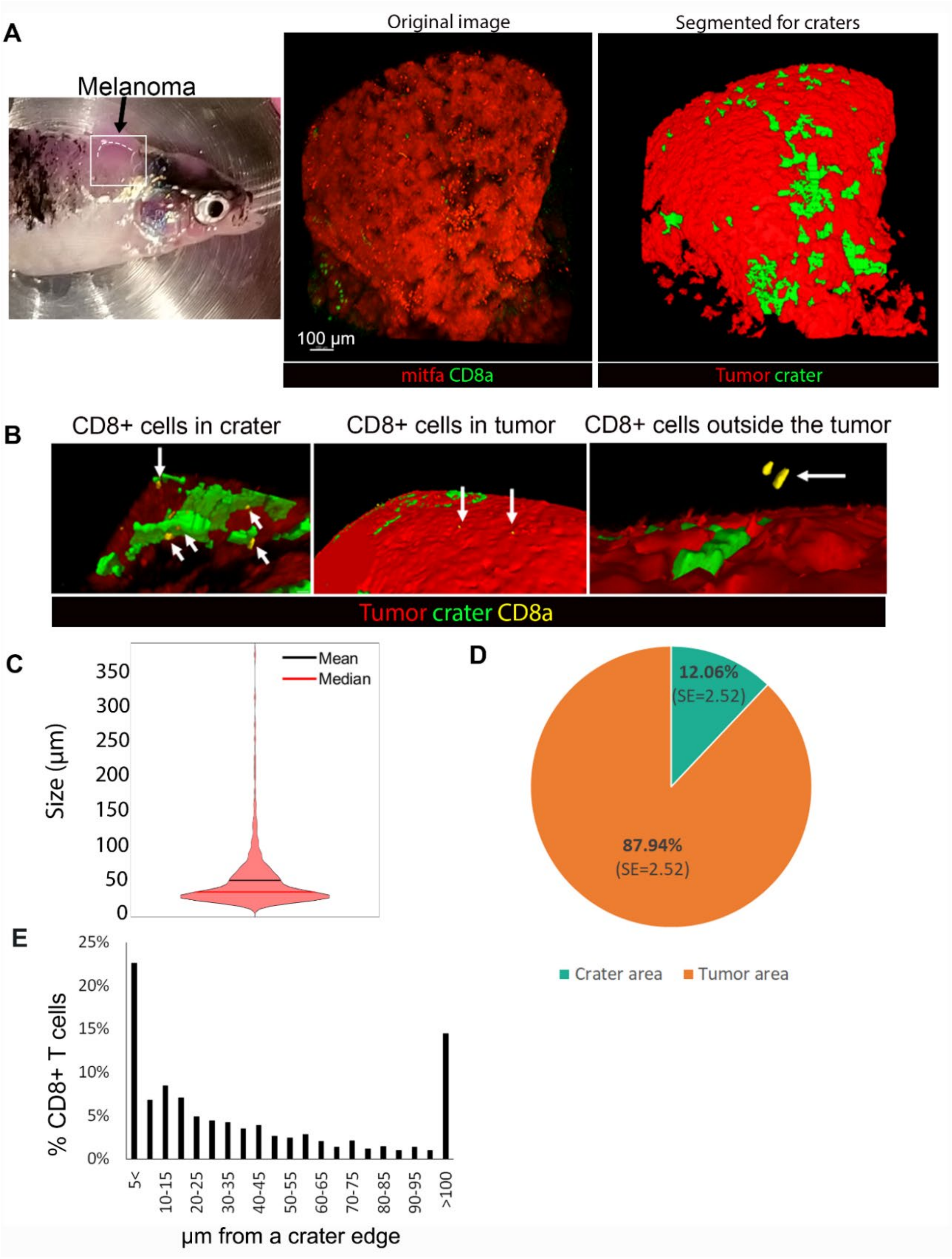


Figure S3. **Quantification of CD8α:EGFP cells-craters interactions. (A**) Left: Picture of a zebrafish with mCherry$^+$ melanoma protruding behind its head (the fish is intubated for long-term time-lapse imaging). Middle: The fluorescent image of the area marked with the white rectangle on the left image. A white dashed line marks the upper border of the confocal imaged tumor area (mirror image). Right: Outcome of tumor and crater segmentation, as described in the materials and methods, of the image in the middle. The areas lacking signal and corresponding to craters are now segmented objects marked in green. (**B**) Examples of CD8$^+$ cells quantified as "in crater" and "in tumor". Yellow arrows mark CD8$^+$ cells. The cells were quantified when directly connecting the segmented areas per segmented surface (mm$^2$). Some CD8$^+$ cells were found in the epithelial layer covering the tumor, i.e. "outside" the tumor, and were not included in the analyses. (**C**) Violin plot of crater size distribution (diameter in µm) in untreated tumors. Craters range mostly from 20µm-100µm, with a mean of 50µm. (n=3 fish/265 craters) (**D**) Crater area constitute about 12% of the tumor surface in untreated tumors (n=5 fish, data is mean$^+$SE). (**E**) CD8$^+$ cell distribution by distance from craters, showing fewer CD8$^+$ cells as the distance from craters increase (n=6 fish, calculated for 1200 cells).

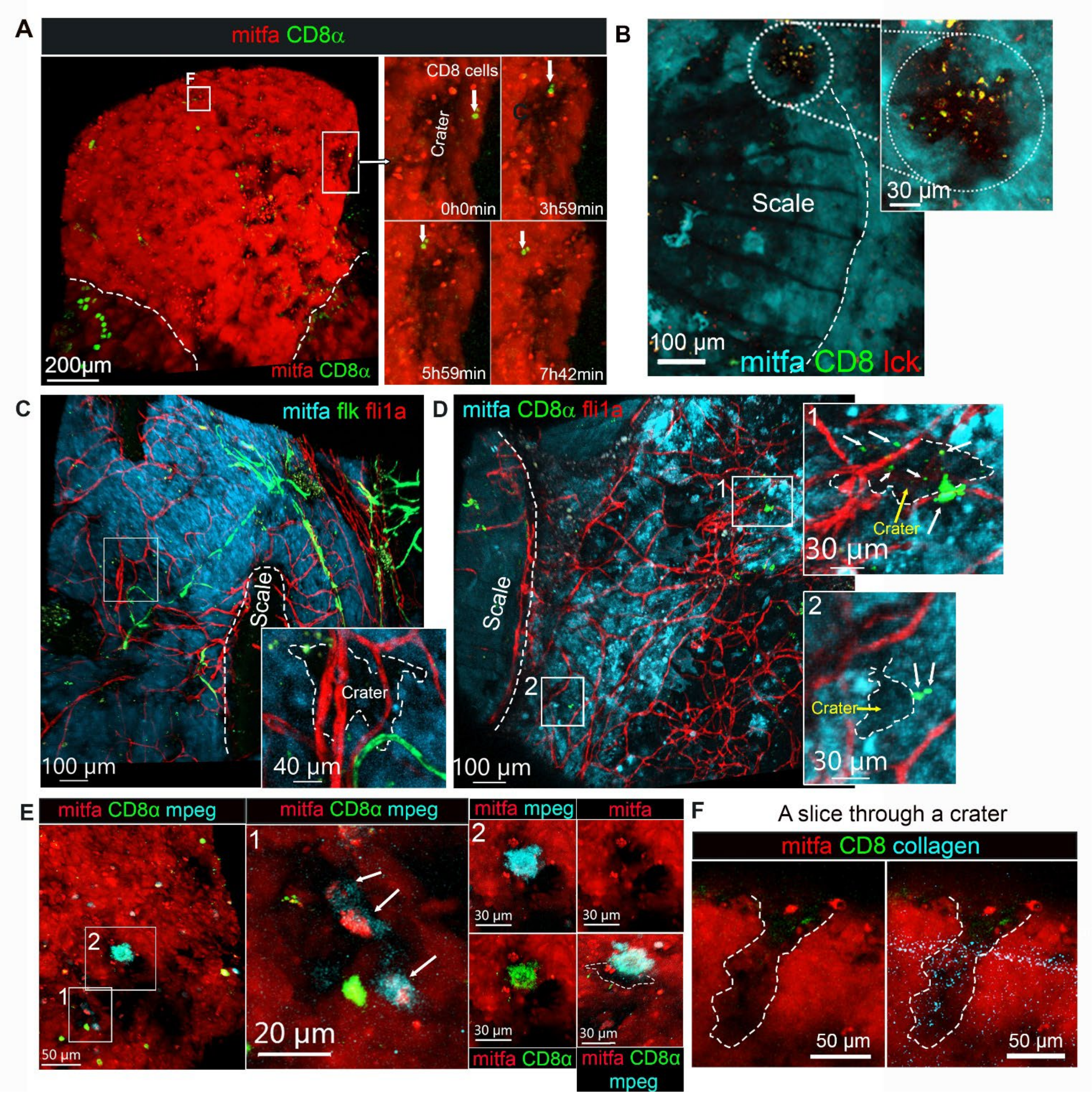


Figure S4. **Craters harbor CD8$^+$ T cells and macrophages and are found near blood vessels.** (**A**) Snapshots of long-term live imaging of the area marked by the large rectangle in the low magnification image to the left. CD8$^+$ T cells linger in a crater for nearly 6 hours before moving out. Time is hours:minutes. (**B**) Crater at the scale edge harboring lck$^+$/CD8α$^+$ T cells. Dashed line marks the scale edge. The crater is circled by dashed line, enlarged to the right. **(C)** 3D image of a tumor in Tg(flk:GFP; fli1a:dsRed) zebrafish. Dashed line outlines a scale protruding from the tumor mass, in the low magnification image. A crater area is enlarged at the low right corner, to show proximity to fli1a$^+$/flk$^-$ and flk$^+$ blood vessels (crater marked by a dashed line). (**D**) 3D image of a tumor in Tg(cd8α:EGFP; fli1a:dsRed) zebrafish. The areas in the rectangles are enlarged to the right. Dashed lines mark the craters. White arrows mark CD8$^+$ T cells. Area (1) shows CD8$^+$ T cells adjacent to blood vessels in a crater. Area (2) shows CD8$^+$ T cells interacting with tumor cells and do not contact blood vessels. The craters emerge adjacent to the fli1a$^+$ blood vessel. (**E**) 3D image of mitfa:mCherry melanoma in Tg(cd8α:EGFP; mpeg1:BFP) zebrafish. Two areas marked by rectangles 1 and 2 are enlarged on the right. Area (1): enlarged view of a crater harboring CD8α$^+$ T cells and mpeg1$^+$ monocytes engulfing melanoma cells (marked by white arrows). Area (2): Large CD8α$^+$/mpeg$^+$ dendritic cell above a crater. **(F)** A slice (2um slice of a z-stack) through the crater marked by the small rectangle F in panel A. Using second harmonics generation, fine collagen fibers can be seen penetrating the crater from the collagen-rich capsule that encompass the tumor (horizontal SHG signal- refer to Figure S7G).

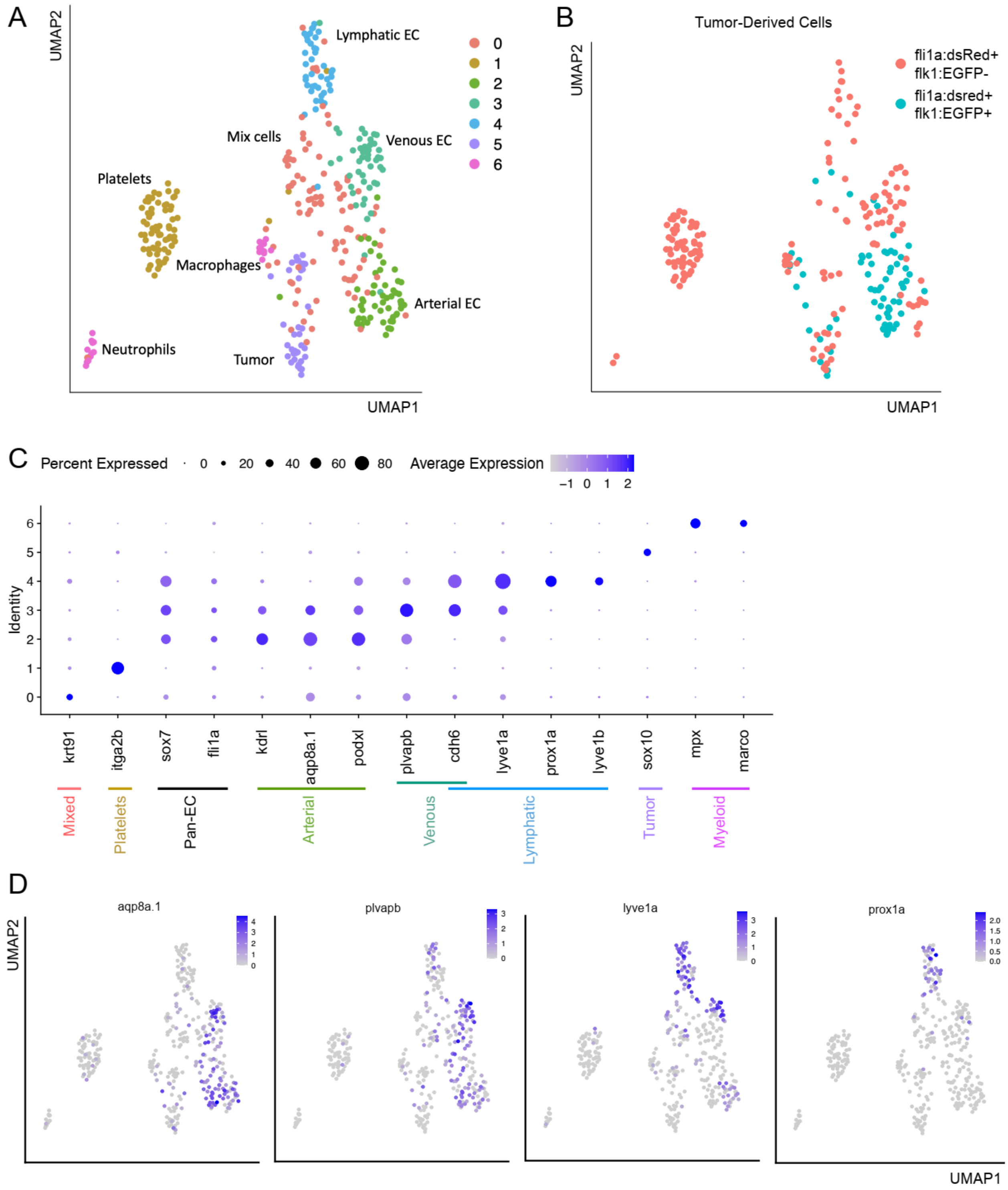


Figure S5. **Flk marks a distinct subset of fli1a$^{+}$ endothelial cells.** (**A**) scRNA-seq analysis of sorted fli1a$^{+}$/flk1$^{+}$ and fli1a$^{+}$/flk1$^{-}$ cells from zebrafish melanoma and normal skin using the SORT-seq technology exhibits 7 clusters. (**B**) fli1a$^{+}$/flk1$^{+}$ cells largely cluster apart from the other tumor-derived cells. (**C**) Differential gene expression analysis identifies 3 endothelial cell clusters, suggestive of arterial (cluster 2), venous (cluster 3), and lymphatic (cluster 4) subpopulations. (**D**) Marker genes exhibit differential expression across endothelial cell subpopulations.

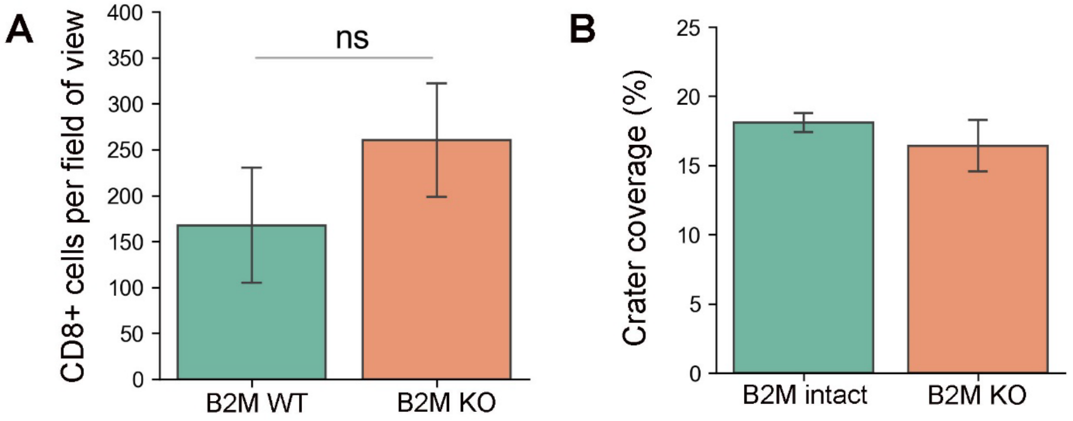


Figure S6. **CD8$^+$ T cells overall number and density within crater in B2M KO tumors.** (**A**) CD8$^+$ cell number per field of view and **(B)** crater coverage (% of the tumor surface), imaged from tumors of intact and B2M-depleted melanomas. (For both graphs: n=4 control fish, 6 field of view, 5 B2M-depleted fish, 8 field of view. Data is mean±SE, T test. p-value for (A)=0.34. p value for (B)=0.12).

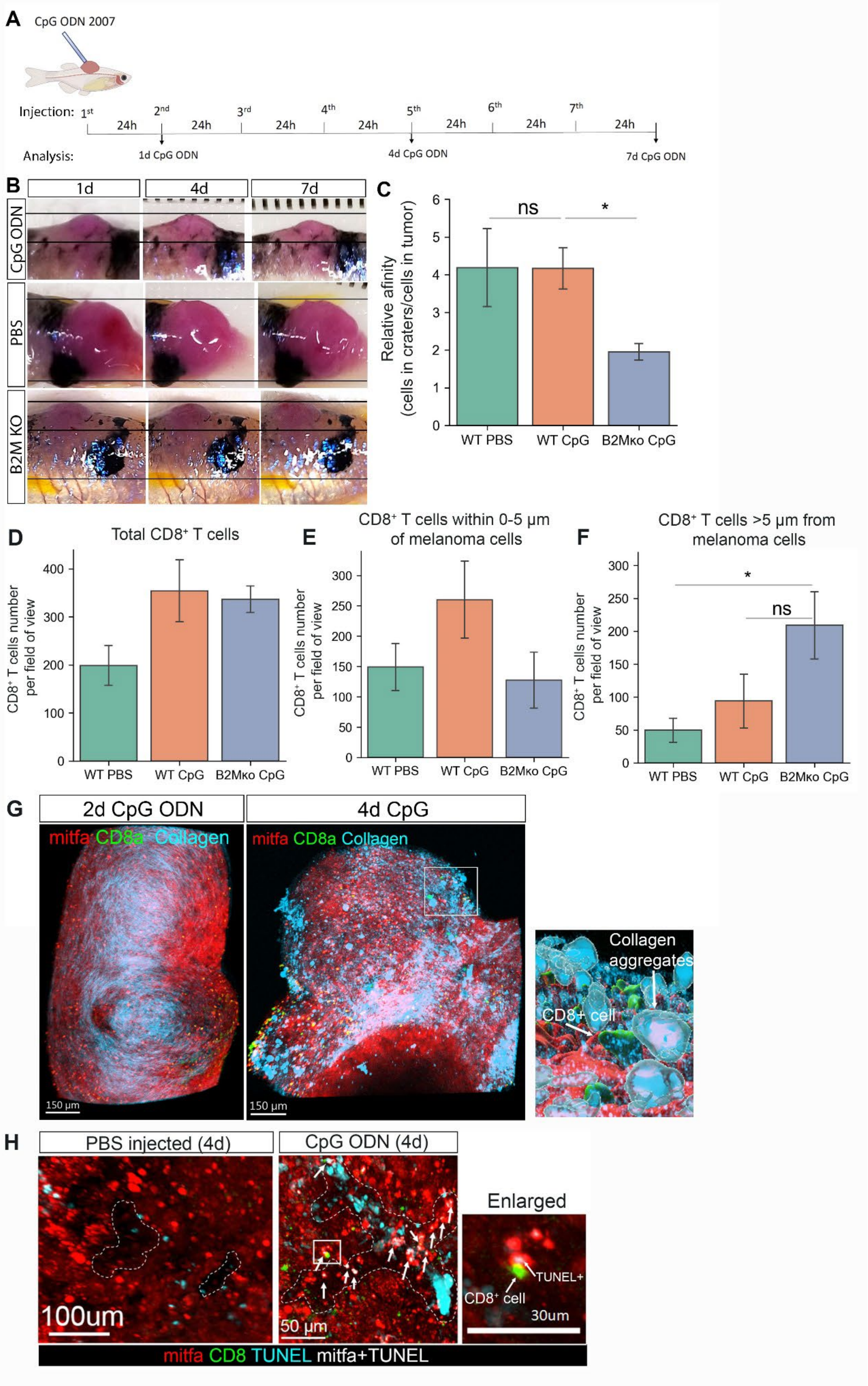




Figure S7. Legend in the next page

Figure S7. **Immune induction by CpG ODN treatment induces B2M dependent tumor shrinkage and cell death.** (**A**) A diagram of CpG ODN treatment applied to zebrafish melanoma. (**B**) Representative images of melanoma tumors injected daily with CpG ODN or PBS for 1,4 and 7 days, each 24 hours post last injection. Upper panel- WT tumor injected with CpG ODN, middle pannel- WT tumor injected with PBS, lower panel B2M KO tumor, injected with CpG ODN (**C**) Relative affinity of $CD8^+$ T cells to craters ($CD8^+$ T cells density in craters/tumor), **(D)** $CD8^+$ T cells numbers per field of view, **(E)** numbers per field of view of $CD8^+$ T cells near or contacting tumor (<5μm away from tumor), and **(F)** numbers per field of view of $CD8^+$ T cells away or not contacting tumor (>5μm away from tumor) in PBS- CpG ODN- and tumor specific B2M KO, CpG ODN- treated melanomas. (calculated for samples presented in Fig. 3B,C: PBS-5, CpG ODN-5, B2M ko CpG ODN-4 fish. 2-3 fields of view per each fish, Data are mean±SE. P value *=0.01). (**G**) 3D image using second harmonic generation (SHG) microscopy to visualize collagen of CpG ODN injected tumors in CD8α:EGFP transgenic zebrafish (representative of 2 CpG ODN 2d- and 4 CpG ODN 4d-injected fish). $CD8^+$ cell infiltration after 4 daily injections is accompanied by multiple collagen aggregations. Area marked by rectangle area is enlarged, tilted for a side view and segmented, showing $CD8^+$ cells in craters along with multiple collagen aggregations **(H)** Whole mount TUNEL assay of PBS- and CpG ODN-injected tumors for 4 days. White dashed lines mark craters' borders at the left and middle images. Arrows indicating $TUNEL^+$ melanoma cells. The area marked by the rectangle on the middle, CpG ODN, image is enlarged on the right, showing a $CD8^+$ cell contacting a $TUNEL^+$ melanoma cell.

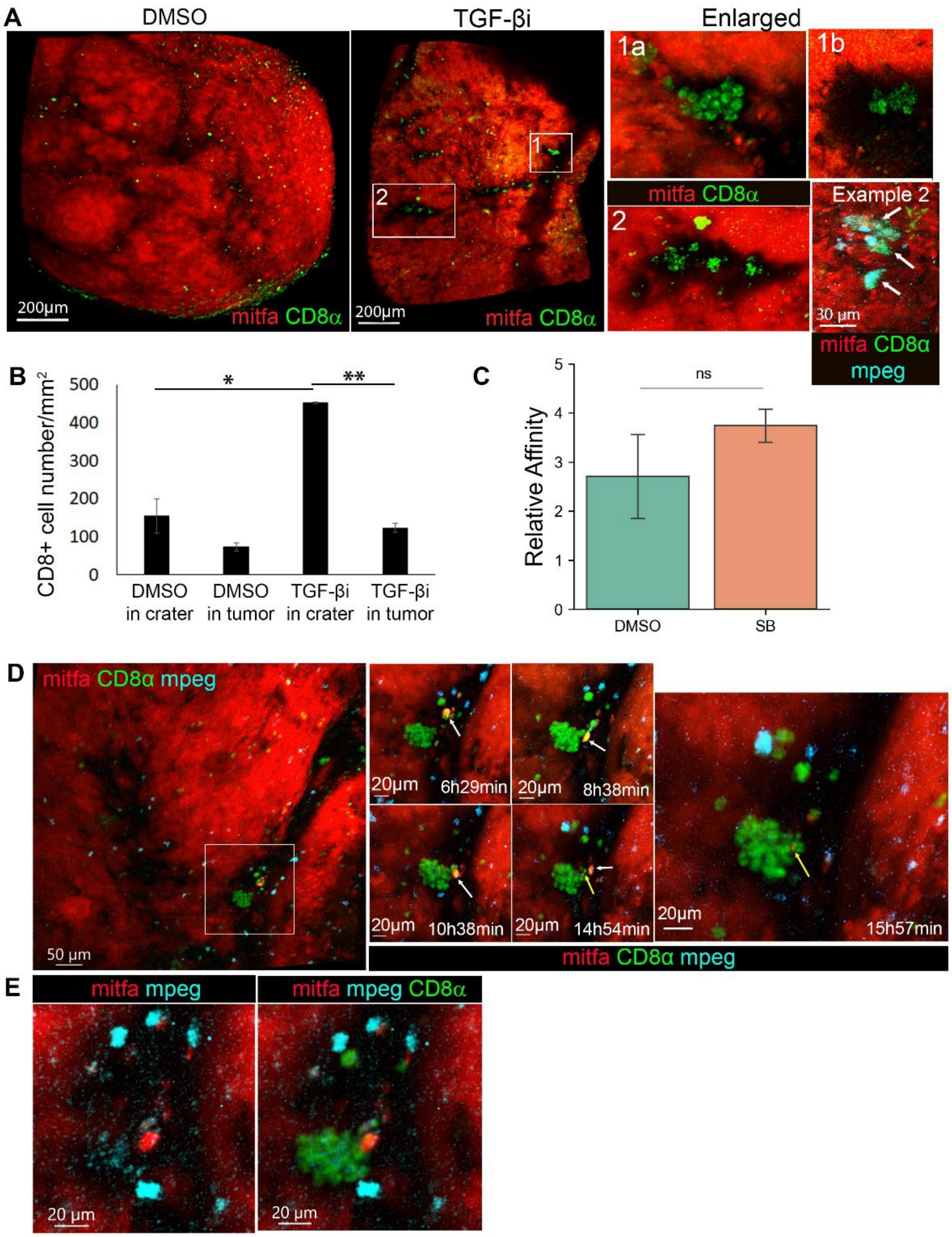


Figure S8. **TGF-β inhibition induces accumulation of CD8⁺ cells within craters.** (**A**) 3D image of melanoma in *cd8a:EGFP* transgenic zebrafish treated with SB431542 (TGF-βi) or DMSO for 24 hours (representative of 3 fish). Enlarged area (1): A $CD8^+$ T cell cluster. (1a): a 3D view of the cluster. (1b): a single slice image at the plane beneath the cluster shown in 1a, revealing a large dendritic-shaped $CD8^+$ cell beneath the cluster. Enlarged area (2): a crater with multiple $CD8^+$ dendritic-shaped cells. Example 2: A different tumor grown on Tg(cd8a:EGFP;mpeg:BFP) zebrafish, treated with SB431542 for 24h, shows a $CD8^+$/$mpeg^+$ myeloid cells (white arrows) cluster in a crater. (**B**) $CD8^+$ cell density in craters vs. tumor in SB431542 treated vs DMSO control. (n=6 DMSO, 3 TGF-βi, TTest. Data is mean±SE ,*p value=0.043, **p value=0.0012). (**C**) Relative affinity of $CD8^+$ cells to craters (cells density in craters/cells in tumor, in TGF-βi vs. DMSO treated fish (n=6 DMSO, 3 TGF-βi, T-Test. Data is mean±SE). (**D**) Snapshots from a 16 hours long-term time-lapse imaging of mitfa:mCherry tumor in Tg(*cd8a:EGFP;mpeg:BFP*) zebrafish following 24h TGF-βi treatment. The area marked by rectangle is enlarged at the right. Over the course of 16 hours, $CD8a^+$ T cells interact with a mCherry melanoma (marked by white arrow) for nearly 9 hours. Then pass it to the $CD8a^+$ dendritic-shaped cell which releases it at hour 15 of imaging, keeping a fragment of mCherry (yellow arrow). Time is hours:minutes.(**E**) Image of the dendritic $CD8a^+$ cell at 9h38min, fluorescent-color separated for mpeg (monocyte marker) and CD8α marker, showing expression of both *mpeg* and *cd8a*.

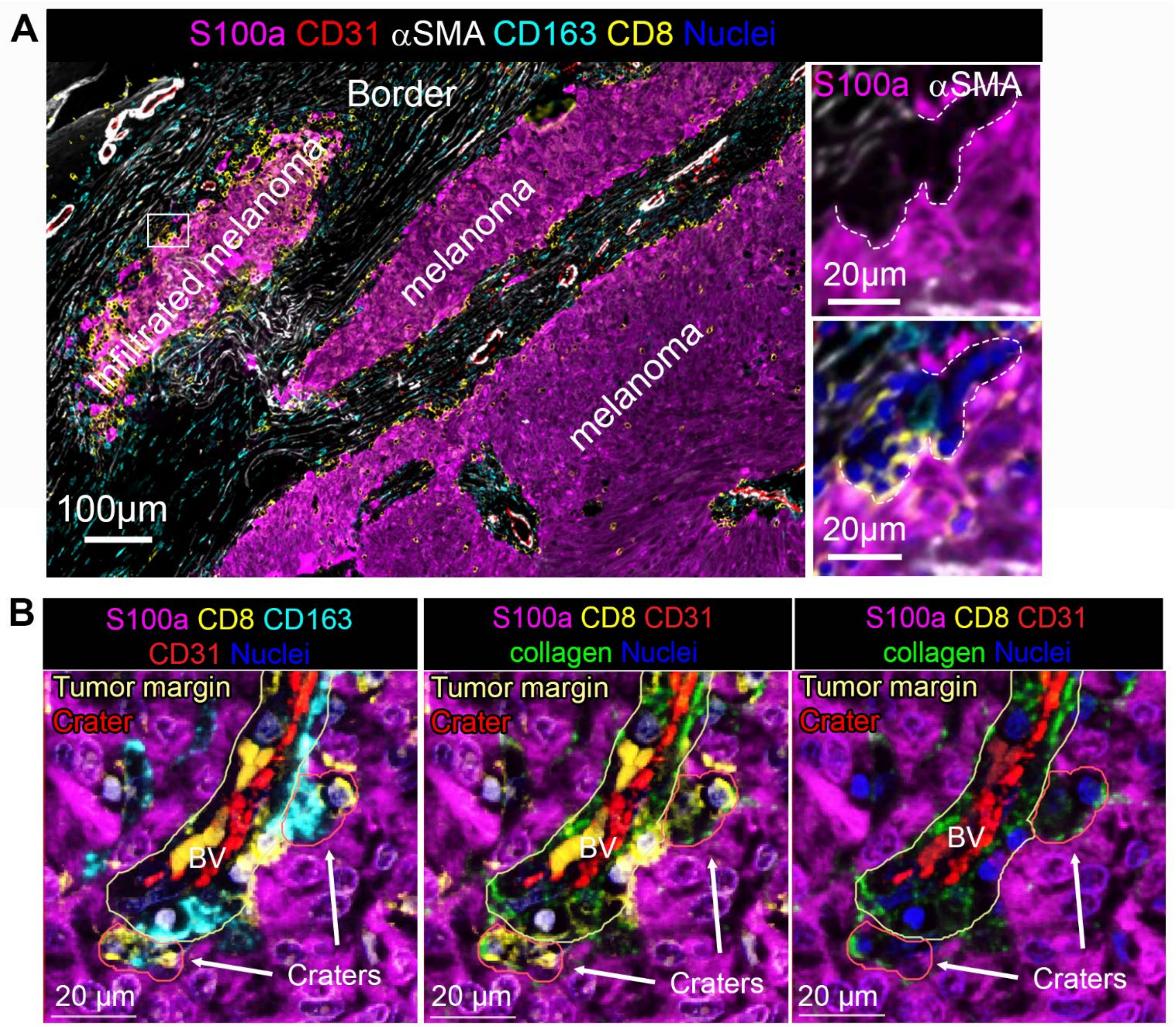


Figure S9. **Craters appear at the tumor border and are lined by collagen fibers protruding from the collagen rich perivascular area. (A)** Left: CyCIF image of $S100^{+}$ melanoma tissue contacting the stromal layer at the border of the tumor. Right: the area marked with rectangle is enlarged, showing a crater containing $CD8^{+}$ T cells and $CD163^{+}$ cell. (**B**) A representative image of collagen lining at craters in an untreated metastatic melanoma. The picture presents a perivascular area (marked by a fine yellow line) with two protruding craters (marked by red lines) containing $CD8^{+}$ T cells and $CD163^{+}$ DCs (left image). The perivascular area is rich with collagen. Collagen fibers are seen extending into the craters (middle and left images).

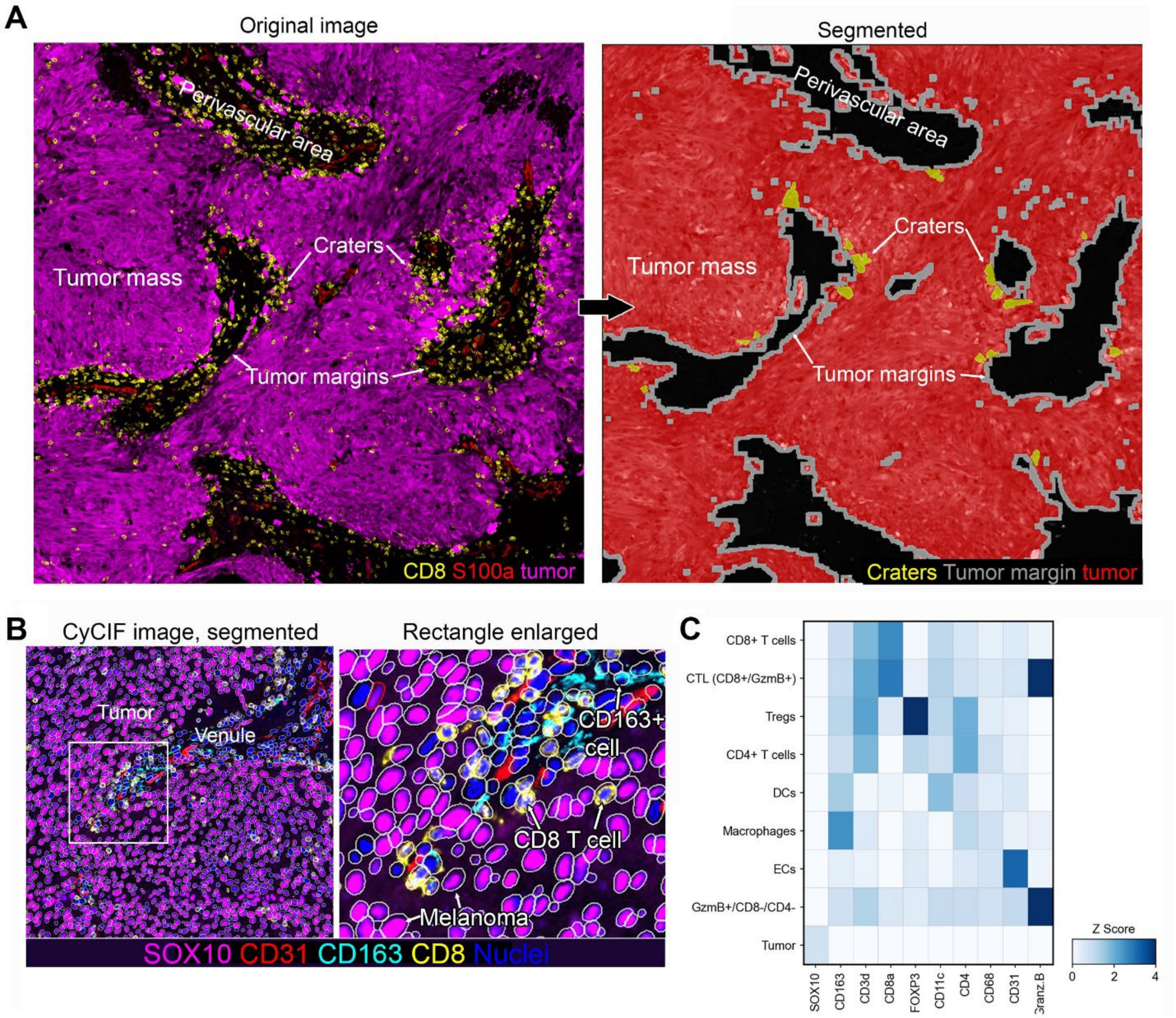


Figure S10. **Quantification analysis of human CyCIF processed untreated melanoma samples.** (**A**) An example of a CyCIF images after segmentation as done in this study. The image on the left is segmented on the right, highlighting the crater areas (yellow), Tumor margins (grey) and Tumor mass (red). See materials and methods for a detailed description of the analysis. **(B)** Example of cell segmentation of a human CyCIF sample of primary human melanoma as analyzed in this study. The area in the white rectangle is enlarged on the right to show the individual cells. Cell segmentation was followed by cell clustering and identification of cell populations stained in the sample: (**C**) Signal intensity heatmap of markers read in the CyCIF samples analysed, identifying cell populations.

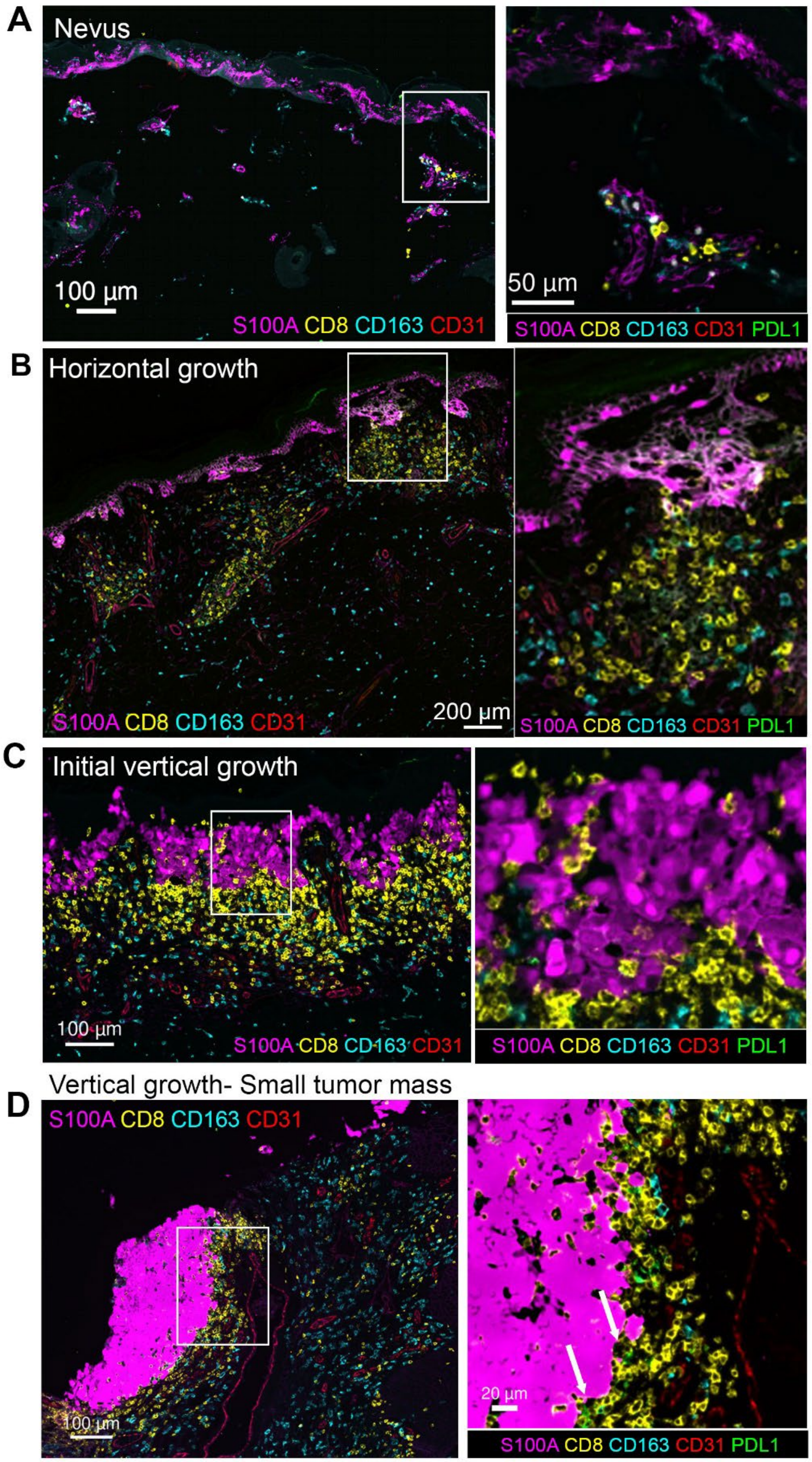


Figure S11. **Craters first appear at the vertical growth phase of melanoma development.** CyCIF images of untreated primary melanoma tumors at different stages of development. For each panel, the area marked by the white rectangle in the left image is enlarged to the right. (**A**) Nevus, with distant $CD8^+$ T cells. (**B**) Horizontal growth, with $CD8^+$ T cells aggregate around blood vessels. (**C**) The tumor thickens and multiple $CD8^+$ T cells contact its margins. PD-L1 negative indentations are seen at the tumor margins. (**D**) Vertical growth area forming a small tumor mass in melanoma at the vertical growth phase. Craters (marked by white arrows) appear at the melanoma border containing $CD8^+$ T cells and PD-L1 expressing $CD163^+$ DCs.

Images are representatives of the following reviewed samples: Superficial radial phase: n=15. Vertical growth: n=3.

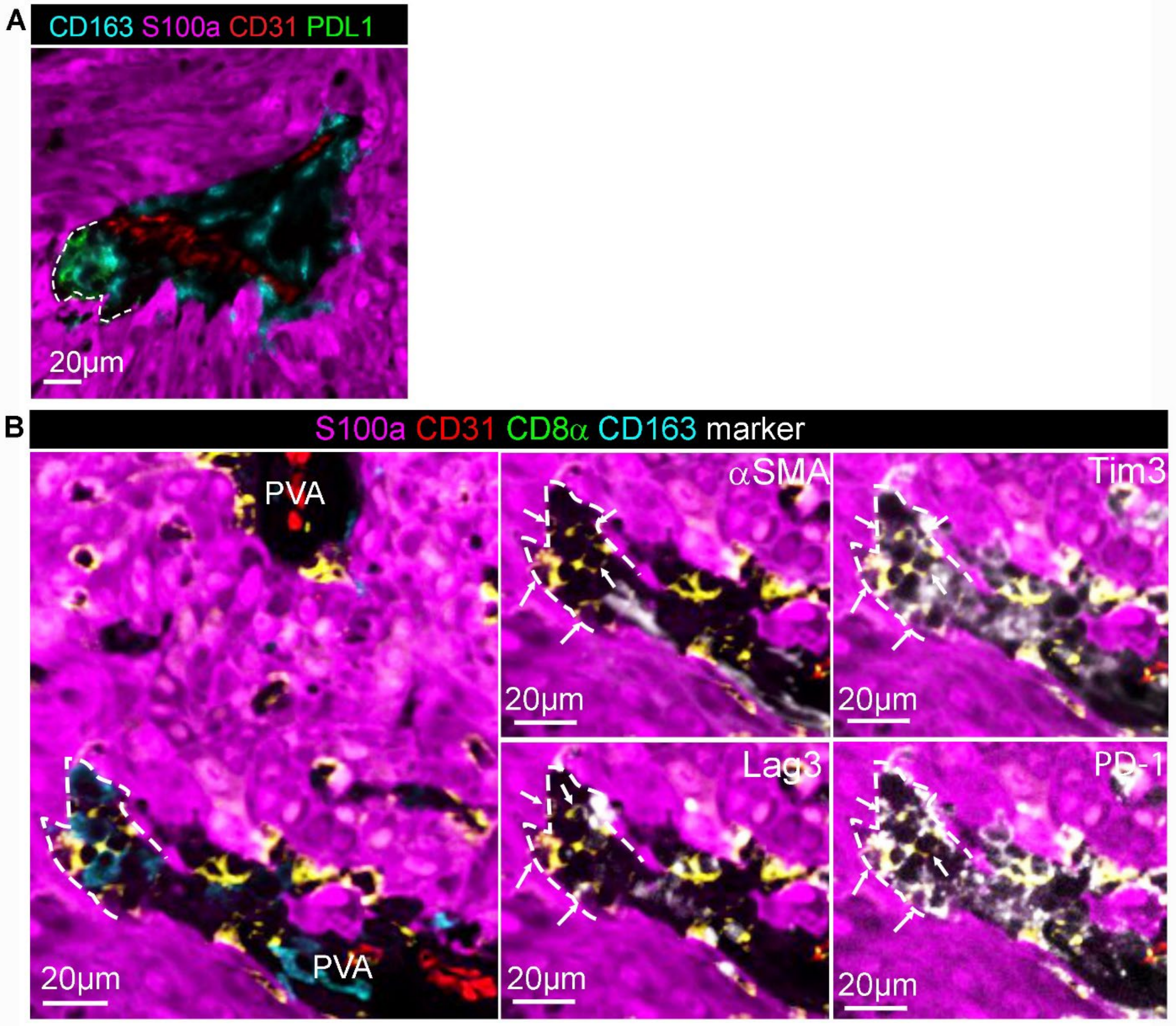


Figure S12. **Human melanoma craters harbor dysfunctional CD8$^+$ T cells.** (**A**) The same crater that is presented in figure 7B, here showing PD-L1 expression on the CD163$^+$ DCs. (**B**) Representative image of a crater (marked by dashed line) breaching out of a perivascular area (PVA). Left: images show that the marked crater. Right: the crater is αSMA negative, containing CD8$^+$ T cells (white arrows) expressing Tim3, Lag3 and PD-1 (representing 3 tumors)

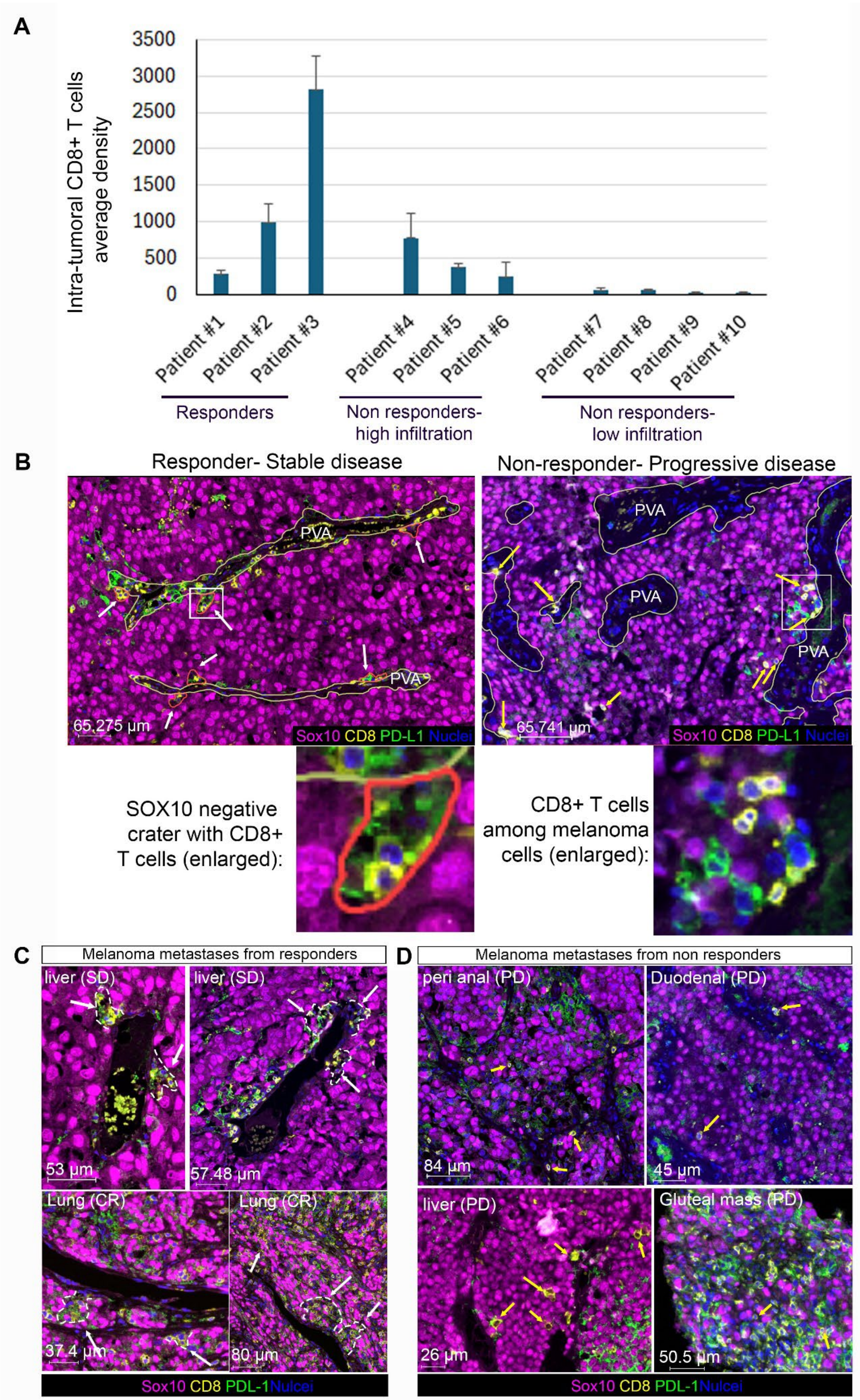


Figure S13. Legend in the next page

Figure S13. **Craters appear in patients responding to ICB, but not in non-responders. (A)** Intra-tumoral $CD8^+$ T cell average density, measured at the pathology department of BWH, of each of the post-treatment samples analyzed. Graph depicts average intra-tumoral $CD8^+$ T cells number per $mm^2$ for at least 6 regions of interest per sample and the corresponding SE. Density below 200 was considered lowly infiltrated. (see materials and methods) **(B)** Representative images of residual melanoma metastases of patients following ICB presenting the different modes of $CD8^+$ T cells infiltration into the tumor mass in responders and non-responders, following ICB treatment. Left: durable response (stable disease). Dashed lines and arrows indicate craters. Right: progressive disease. Yellow arrows indicate $CD8^+$ T cells entering the tumor. At the bottom of each image: enlarged view of the area marked by white rectangles in the images above. While in durable response, $CD8^+$ T cells aggregate in craters (craters marked by a dashed line), in progressive disease most $CD8^+$ T cells are found as singles or doublets (yellow arrows). Even when $CD8^+$ T cells clustered together at the stroma-tumor border (enlarged area), they were not found in craters but were spread among melanoma cells. **(C)** Additional Representative images reviewed samples of melanoma residual metastasis of patients presenting a durable response to ICB treatment. Upper: two additional areas from the stable disease sample- the same sample shown in Figure 6E. Lower: samples from two patients who presented complete response clinically. Each image represents a different sample. Craters are marked by white dashed line and arrows in patient responding to ICB. (**C**) Four representative samples out of seven analyzed samples of melanoma residual metastasis from patients who did not respond to ICB and presented clinically a progressing disease. Each image represents a different sample. Yellow

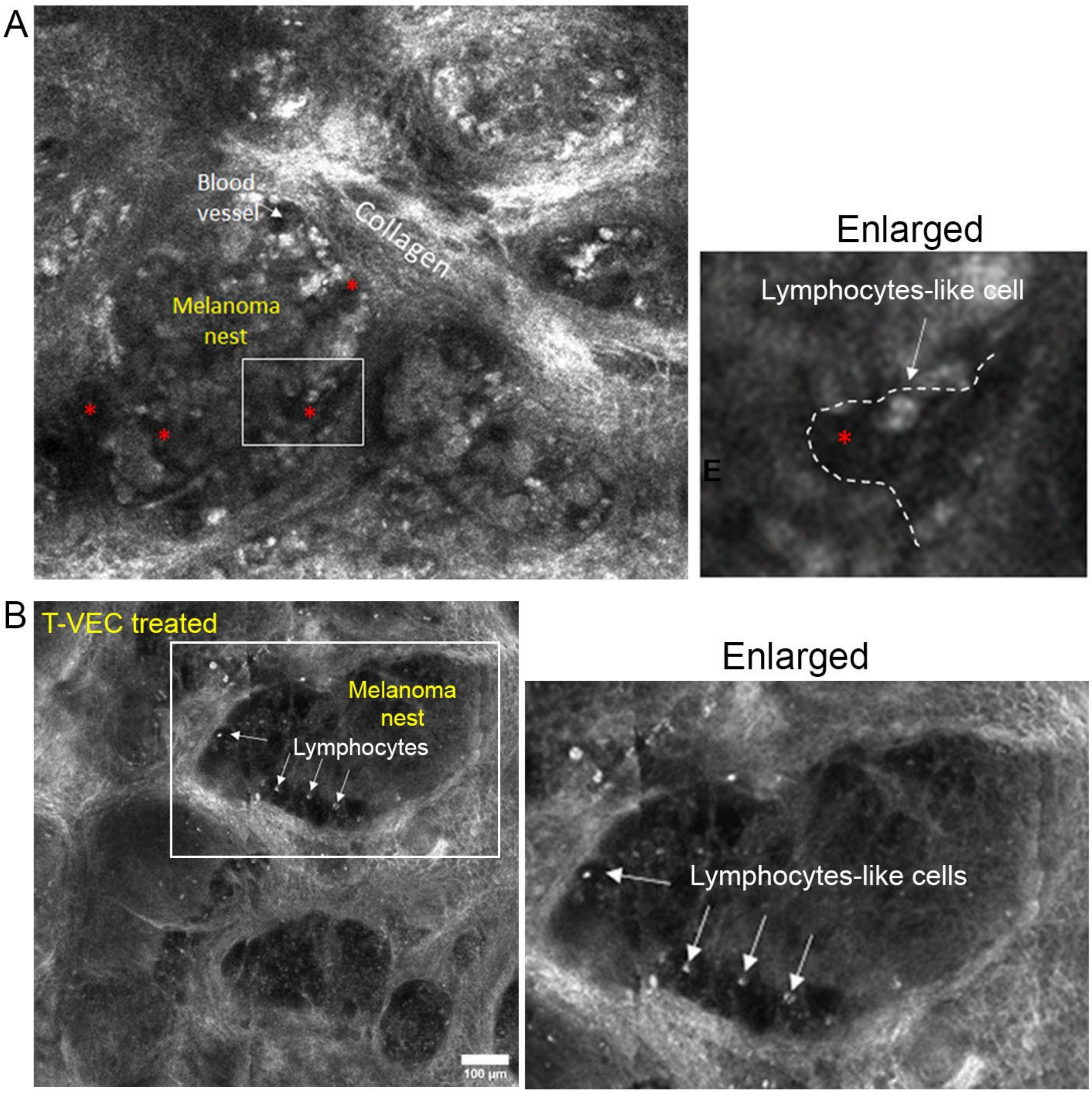


Figure S14. **Confocal imaging of human melanoma tumors in vivo reveal crater like structures**. (**A**) in vivo, confocal view of melanoma nests in patients using RCM. Craters can be found at the edge of the nest (enlarged). (**B**) Melanoma nests in a patient treated with T-VEC. The nests are emptying of cells and large craters appear together with lymphocyte-like cells (enlarged). White arrows indicate lymphocyte like cells, as determined by pathologists reviewing the images (see materials and methods).

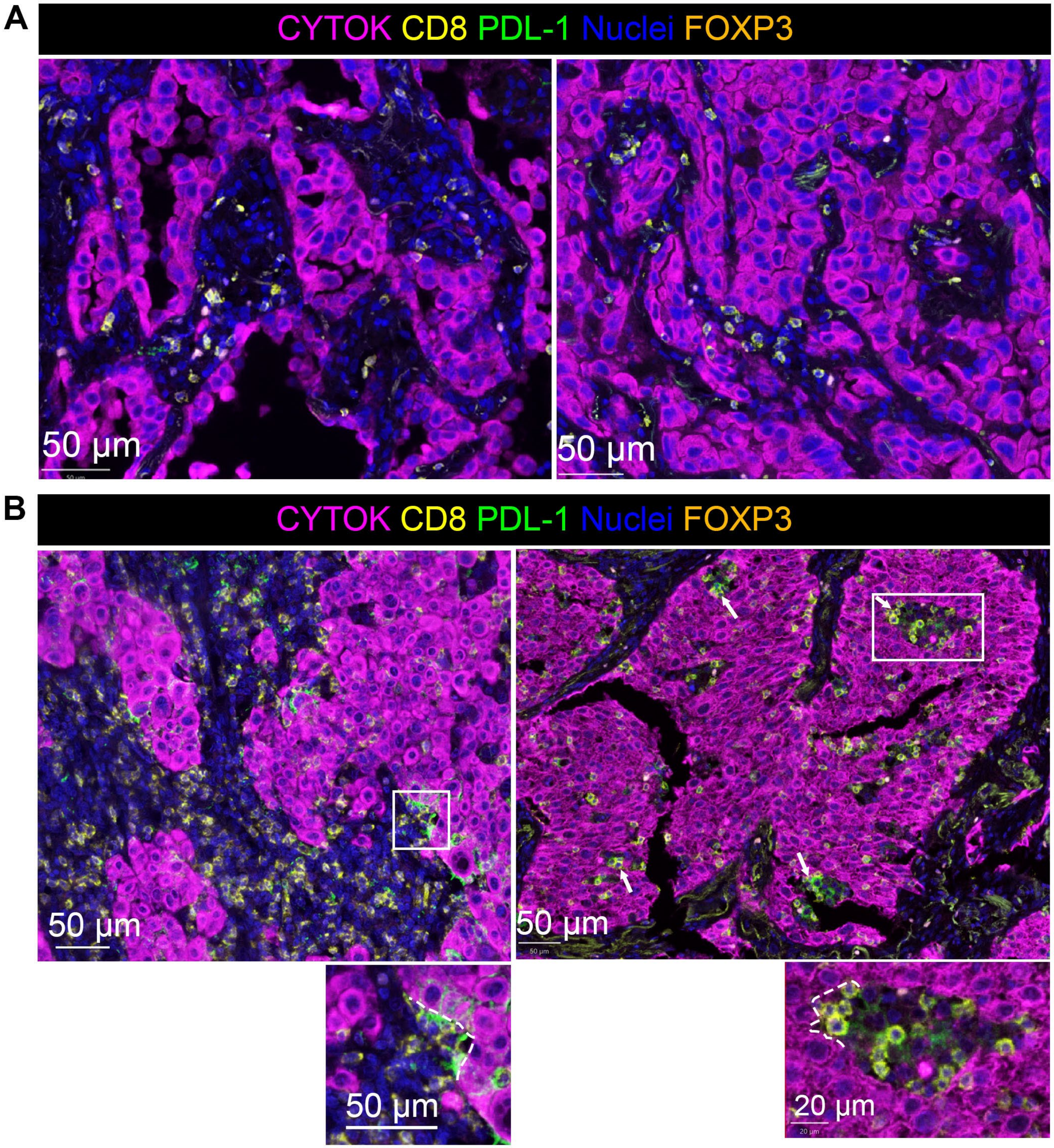


Figure S15. **Craters appear in non-small cell lung cancer (NSCLC) in a developed tumor mass.** (**A**) Two representative images of NSCLC not yet forming a lobular mass. $CD8^+$ T cells are mostly scattered at the stromal area; single cells engage tumor cells. (**B**) Two representative images of developed tumors. Multiple $CD8^+$ T cells at the stromal area. $CD8^+$ T cells clusters appear in pockets breaching into $CYTOK^+$ lung cancer tumor mass. White arrows point at craters at the right image. Areas marked by white rectangles are enlarged at the bottom of each image, where crater areas are marked by white dashed lines. Representatives of 15 samples reviewed.

Table S1. List of multiplex immunofluorescence processed human melanoma samples reviewed in the paper

| | Tumor type | Tumor site | pre/post ICB treatme | ICB respns | Infiltration level |
|---|---|---|---|---|---|
| #1 | Primary melanoma | Skin | pre treatment | N/A | low infiltration |
| #2 | Primary melanoma | Skin | pre treatment | N/A | low infiltration |
| #3 | Primary melanoma | Skin | pre treatment | N/A | low infiltration |
| #4 | Primary melanoma | Skin | pre treatment | N/A | low infiltration |
| #5 | Primary melanoma | Skin | pre treatment | N/A | low infiltration |
| #6 | Primary melanoma | Skin | pre treatment | N/A | low infiltration |
| #7 | Primary melanoma | Skin | pre treatment | N/A | low infiltration |
| #8 | Primary melanoma | Skin | pre treatment | N/A | low infiltration |
| #9 | Primary melanoma | Skin | pre treatment | N/A | high infiltration |
| #10 | Primary melanoma | Skin | pre treatment | N/A | high infiltration |
| #11 | Primary melanoma | Skin | pre treatment | N/A | high infiltration |
| #12 | Primary melanoma | Skin | pre treatment | N/A | high infiltration |
| #13 | Primary melanoma | Skin | pre treatment | N/A | high infiltration |
| #14 | Primary melanoma | Skin | pre treatment | N/A | high infiltration |
| #15 | Primary melanoma | Skin | pre treatment | N/A | high infiltration |
| #16 | Melanoma metastasis | Soft tissue | pre treatment | N/A | low infiltration |
| #17 | Melanoma metastasis | Bowel | pre treatment | N/A | low infiltration |
| #18 | Melanoma metastasis | Bowel | pre treatment | N/A | low infiltration |
| #19 | Melanoma metastasis | Lung | pre treatment | N/A | low infiltration |
| #20 | Melanoma metastasis | Skin | pre treatment | N/A | low infiltration |
| #21 | Melanoma metastasis | Skin | pre treatment | N/A | low infiltration |
| #22 | Melanoma metastasis | Skin | pre treatment | N/A | low infiltration |
| #23 | Melanoma metastasis | Soft tissue | pre treatment | N/A | low infiltration |
| #24 | Melanoma metastasis | CNS/Brain | pre treatment | N/A | low infiltration |
| #25 | Melanoma metastasis | Liver | pre treatment | N/A | high infiltration |
| #26 | Melanoma metastasis | Lung | pre treatment | N/A | high infiltration |
| #27 | Melanoma metastasis | Lung | pre treatment | N/A | high infiltration |
| #28 | Melanoma metastasis | Bowl | pre treatment | N/A | high infiltration |
| #29 | Melanoma metastasis | Bowl | pre treatment | N/A | high infiltration |
| #30 | Melanoma metastasis | Bowl | pre treatment | N/A | high infiltration |
| #31 | Melanoma metastasis | Bowl | pre treatment | N/A | high infiltration |
| #32 | Melanoma metastasis | Soft tissue | pre treatment | N/A | high infiltration |
| #33 | Melanoma metastasis | Soft tissue | pre treatment | N/A | high infiltration |
| #34 | Melanoma metastasis | Lung | pre treatment | PR | Low infitration |
| #35 | Melanoma metastasis | Lung | pre treatment | PD | Low infitration |
| #36 | Melanoma metastasis | Pubic ramus | pre treatment | CR | High infilrtation |
| #37 | Melanoma metastasis | Skin | pre treatment | PR | High infilrtation |
| #38 | Melanoma metastasis | Skin | pre treatment | PR | Low infitration |
| #39 | Melanoma metastasis | Chest wall | pre treatment | CR | Low infitration |
| #40 | Melanoma metastasis | Skin | Post treatment | PR | Low infiltration |
| #41 | Melanoma metastasis | Lung | Post treatment | CR | High infiltration |
| #42 | Melanoma metastasis | Around trachea | Post treatment | SD | Low infiltration |
| #43 | Melanoma metastasis | Skin | Post treatment | PD | Low infiltration |
| #44 | Melanoma metastasis | Liver | Post treatment | PD | Low infiltration |
| #45 | Melanoma metastasis | Duodenal mass | Post treatment | PD | Low infiltration |
| #46 | Melanoma metastasis | Skin | Post treatment | PD | High infiltration |
| #47 | Melanoma metastasis | Liver | Post treatment | PD | Low infiltration |
| #48 | Melanoma metastasis | Colon | Post treatment | PD | Low infiltration |
| #49 | Melanoma metastasis | Anal | Post treatment | PD | Low infiltration |

ICB response abbreviations: CR-complete response. PR-partial response. SD-stable disease. PD-progressive disease. Does not include CyCIF data.

Table S2. Antibodies used for multiplex immunofluorescence of human melanoma samples.

| Antibody | Clone | Company | Catalog # | Antibody Dilution | Fluor | Fluor Dilution | Antigen Retrieval |
|---|---|---|---|---|---|---|---|
| COLLAGEN | EPR7785 | Abcam | ab138492 | 1:100 | Opal 570 | 1:100 | ER 1, 10 min |
| SOX10 | EP268 | Cell Marque | AC-0237A | 1:100 | Opal 780<br>TSA-DIG | 1:25<br>1:50 | ER 1, 10 min |
| CD31 | ab28364 | Abcam | ab28364 | 1:100 | Opal 520 | 1:100 | ER 1, 20 min |
| CD8 | C8/144B | Dako | M710301-2 | 1:100 | Opal 480 | 1:100 | ER 1, 20 min |
| S100a | EPR525 | Abcam | ab133519 | 1:1200 | Opal 690 | 1:200 | ER1, 10 min |
| CD163 | 10D6 | Leica | CD163-L-CE | 1:100 | Opal 620 | 1:100 | ER1, 20 min |

**Supplementary videos legend**

Video S1. **A CD8$^+$ T cell interacts with a melanoma cell in a crater in untreated melanoma tumor.** Long-term, time-lapse in vivo imaging of the CD8$^+$ T cell (green) in crater in a mitfa:mCherry tumor (red) grown in a cd8a:EGFP transgenic fish, depicted in Fig. 1D. The CD8$^+$ T cell forms a lengthy interaction with a melanoma cell for 5 hours, then enters the crater. Time depicts hh:mm:ss.ms from the beginning of the imaging.

Video S2. **CD8$^+$ T cells linger in a crater in untreated melanoma tumor.** Long-term, time-lapse in-vivo imaging of a crater in mitfa:mCherry tumor (red) grown in a cd8a:EGFP transgenic fish, depicted in Fig. S4A. Two CD8$^+$ cells (green) are shown interacting with melanoma cells at the walls of the crater and moving within it for nearly 6 hours. Time depicts hh:mm:ss.ms from the beginning of the imaging.

Video S3. **CD8$^+$ T cells interact with melanoma cell in a manner characteristic to tumor killing in vivo.** Long-term, time-lapse in vivo imaging of CD8$^+$ T cell cluster interacting with a melanoma cell in a BFP expressing tumor in a (cd8a:EGFP;lck:mCherry) transgenic fish depicted in Fig. 1E. A crater at the scale edge contains CD8$^+$/lck$^+$ T cells (yellow, due to green and red mix) interacting with a centered BFP$^+$ melanoma cell (blue) for over 2 hours. CD8$^+$ T cells can be seen arriving to the melanoma cell from the edge of the crater during this time. Time depicts hh:mm:ss.ms from the beginning of the imaging.

Video S4. **CD8$^+$ T cells interact with melanoma cells in enlarged craters following CpG ODN treatment.** Long-term, time-lapse in-vivo imaging of a mitfa:mCherry tumor in (cd8a:EGFP;mpeg:BFP) transgenic fish injected daily with CpG ODN for 4 days and imaged 24 hours after last injection. The crater border is marked with a dashed line at the beginning of the movie. Multiple CD8$^+$ T cells (green) interact with melanoma cells (red) in the craters. Blue cells are mpeg$^+$ macrophages. Time depicts hh:mm:ss.ms from the beginning of the imaging.

Video S5. **mCherry fragment uptake by CD8$^+$ dendritic cell following TGF-β inhibition.** Long-term, time-lapse in-vivo imaging of mCherry expressing tumor in (cd8a:EGFP;mpeg:BFP) transgenic fish, after immersion with TGF-β inhibitor SB431542 for 24 hours, depicted in Fig. S8D. A crater at the edge of the scale (marked by white dashed line at the beginning of the movie) harbors a cluster of CD8$^+$ T cells (green, small) interacting with a melanoma cell (red) for 10 hours, then transferring it to a CD8$^+$ dendritic cells (green, large). The CD8$^+$ DC interacts with the melanoma cell for 8 hours and then releases it, leaving a fragment of mCherry$^+$ material behind in the CD8$^+$ DC. Time depicts hh:mm:ss.ms from the beginning of the imaging.

Video S6. **3D projection of human melanoma tumor nest, containing a crater-like structure, in a live patient.** 3D projection of reflective confocal microscopy imaging of a melanoma nest in vivo in a patient. An area of irregular borders devoid of cells resembling a crater is marked by yellow dashed line. A small bright cell identified as lymphocyte-like is found at its border.
